# The molecular triggers of human labour: a longitudinal plasma proteomics study

**DOI:** 10.64898/2026.07.07.736770

**Authors:** Katerina Nastou, Nikolai Madrid Scheller, Matteo Tiberti, Anders Tolver, Marie-Louise H. Rasmussen, Elena Papaleo, Stephen Quake, Heather A. Boyd, Mads Melbye

**Affiliations:** Department of Congenital Disorders, Statens Serum Institut, Copenhagen, Denmark; Department of Surgical Gastroenterology, Copenhagen University Hospital Rigshospitalet, Denmark; Danish Cancer Institute, Copenhagen, Denmark; Department of Health Technology, Technical University of Denmark, Denmark; Department of Bioengineering, Stanford University, USA; Department of Applied Physics, Stanford University, USA; Department of Pediatrics, Stanford University School of Medicine, Stanford, California; HUNT Center for Molecular and Clinical Epidemiology, Norwegian University of Science and Technology (NTNU), Trondheim, Norway

## Abstract

Spontaneous labour onset is a precisely timed physiological transition that determines outcomes for millions of pregnancies annually, yet its upstream molecular triggers remain unknown. Here we report a longitudinal plasma proteomics study using the Olink Explore HT platform to measure over 5,400 protein targets in repeated blood samples (median 13 samples per woman) taken from 40 women in the last month up to spontaneous term labour and delivery. Combining longitudinal trajectory modelling with interpretable machine learning for labour timing prediction, we identified five proteins – AFP, ACTA2, ANGPT2, IL1RL1, and LMOD1 – that change in a reproducible sequential order preceding labour onset across all 40 women. IL1RL1, encoding both soluble and membrane-bound ST2, the receptors for IL-33, rose earliest and most consistently, preceding declining AFP and ANGPT2 and a late rise in the smooth muscle contractile proteins ACTA2 and LMOD1. The temporal ordering of these proteins is consistent with a two-phase biological cascade in which feto-placental maturation and vascular remodelling precede activation of the IL-33/ST2 alarmin axis, until the soluble/membrane bound ST2 (sST2/ST2L) balance shifts towards membrane-bound signaling, driving myometrial contractile priming. Mendelian randomisation provided independent human genetic support for the involvement of the IL-33/ST2 axis in labour timing. These findings provide longitudinal plasma proteomic evidence implicating activation of this axis in the timing of spontaneous human labour and identify LMOD1 as a novel circulating marker of myometrial activation.

## Introduction

Globally, approximately 130 to 140 million children are born every year. Thus, onset of labour is one of the most precisely timed physiological transitions in human biology, yet the molecular signals that determine when it occurs remain poorly understood. Labour is currently recognized as a coordinated inflammatory process initiated by endogenous danger signals released from senescent or stressed cells from multiple reproductive tissues (the myometrium, cervix, decidua, and fetal membranes)^1–5^. However, the upstream molecular signals that initiate this inflammatory transition, the order in which inflammatory and contractile events are activated, and the proteins that mark each phase in the systemic circulation, remain largely unknown.

Plasma proteomics provide a window into the dynamic molecular landscape of pregnancy by quantifying thousands of circulating proteins originating from multiple tissues and biological systems. When combined with high-frequency longitudinal sampling, this approach enables detailed mapping of the molecular transitions that precede labour onset. Previous investigations^6–8^ have largely lacked the temporal resolution required to resolve these dynamic changes, leaving key regulatory mechanisms incompletely understood. Untargeted proteomic profiling with dense pre-labour sampling may reveal protein trajectories and signaling pathways not captured by previous approaches, offering new insights into the molecular events behind the initiation of parturition.

Here, we present longitudinal plasma proteomic profiling of 40 women with spontaneous term labour, leveraging dense sampling across the last month preceding labour to define plasma protein trajectories associated with labour timing. Combining longitudinal modelling and interpretable machine learning, we identify a reproducible proteomic signature linking feto-placental maturation, vascular remodelling, inflammatory alarmin signaling, and smooth muscle activation, converging on a sequential biological programme that precedes labour onset with remarkable consistency across individuals. These signals implicate the IL-33/ST2 alarmin axis, a previously unappreciated component of the labour timing mechanism, and identify myometrial activation as a late but consistent downstream consequence of the activation of this pathway. Together, these findings advance our understanding of the molecular mechanisms underlying parturition and open new avenues for investigating the biological programmes that govern the timing of human birth.

## Results

### Data Collection

This study used a group of 40 healthy pregnant women drawn from the Biological Signals in Pregnancy (BSG) Study cohort (see Methods). 491 serial maternal plasma samples from 40 pregnant women (range: 5–15 samples per woman, median: 13) were obtained during the last month of pregnancy at multiple gestational timepoints, with an additional sample collected postpartum (typically 2–12 weeks after delivery). All samples were processed according to standard biobanking protocols^9^ and stored at –80°C until analysis. **Figure 1** shows the sampling days relative to the day of delivery. The 40 women were divided into an exploratory group (n=30) and a confirmatory group (n=10) using k-means clustering on maternal age at delivery, gestational age at delivery, and parity, ensuring representative coverage of clinical covariates in both groups (**Table 1**). To confirm that our population was representative of the general population with respect to gestational age, we compared its gestational age distribution with that of the Early Pregnancy Study^10^. Results of this comparison are presented in **Supplementary Note 1**.

**Figure 1.**
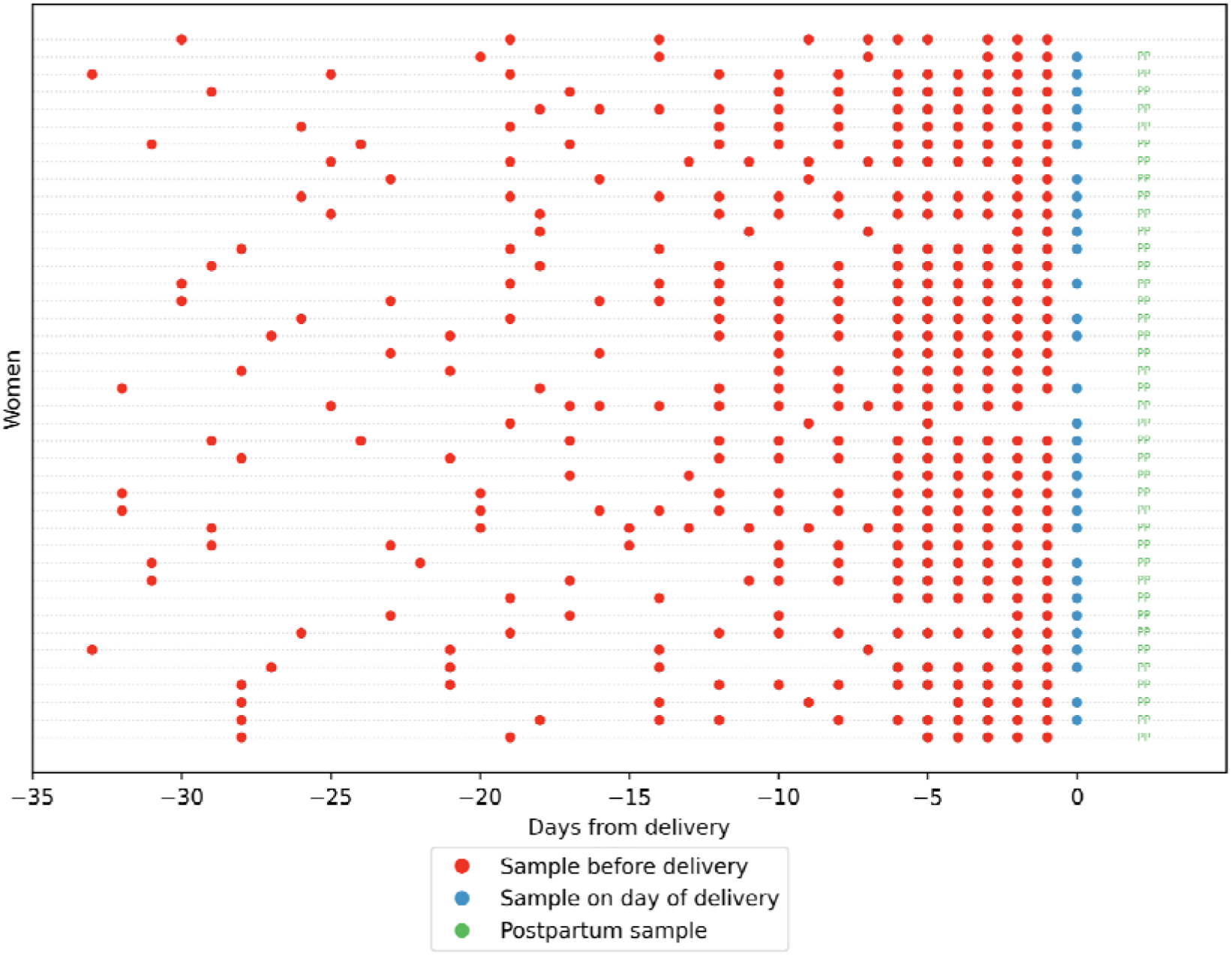
Sample collection timeline relative to delivery. Each dot is one blood sample from a single participant (rows correspond to individual women). The x-axis shows days from delivery: negative values are antepartum samples, 0 is the day of delivery. Colours indicate timing: red = before delivery, blue = on the day of delivery, green = after delivery. Multiple dots along a row indicate repeated sampling for the same woman. “PP” indicates that a postpartum sample exists for that woman. The range of time at postpartum sampling was 7 to 289 days after delivery.

**Table 1.** Demographics and birth characteristics of the exploratory and confirmatory groups. Continuous variables are presented as mean ± standard deviation (SD), and categorical variables are presented as number (percentage). P-values compare the distribution of each variable in the exploratory and confirmatory groups. For continuous variables, p-values were calculated using the Welch two-sample t-test. For categorical variables, p-values were calculated using Pearson’s chi-squared test or Fisher’s exact test when expected cell counts were less than 5. For multi-level categorical variables, the p-value refers to the overall comparison across categories. Maternal age was the age when the first sample was taken, and maternal weight refers to pre-pregnancy weight.

|  | Exploratory group<br>(N=30) | Confirmatory<br>group (N=10) | p-value |
| --- | --- | --- | --- |
| Maternal age at first sample, years | 30.6 ± 2.9 | 31.3 ± 2.9 | 0.53 |
| Height, cm | 167.5 ± 5.7 | 169.9 ± 6.4 | 0.31 |
| Weight, kg | 61.5 ± 8.6 | 60.7 ± 6.4 | 0.76 |
| BMI, kg/m<sup>2</sup> | 22.0 ± 3.4 | 21.0 ± 1.5 | 0.23 |
| Number of previous live births | 0.5 ± 0.7 | 0.4 ± 0.5 | 0.69 |
| Parity, No. (%) |  |  | 0.63 |
| 0 | 19 (63.3) | 6 (60.0) |  |
| 1 | 8 (26.7) | 4 (40.0) |  |
| $\geq 2$ | 3 (10.0) | 0 (0.0) | |
| Mode of delivery |  |  | 1.00 |
| Vaginal, No. (%) | 28 (93.3) | 9 (90.0) |  |
| Cesarean, No. (%) | 2 (6.7) | 1 (10.0) |  |

### A proteome-wide longitudinal screen identifies proteins tracking labour proximity

Preliminary data exploration using principal components analysis (PCA) revealed that the first two principal components clearly delineated whether a sample was obtained antepartum or postpartum; in uniform manifold approximation and projection (UMAP) projections, the overall data structure was dominated by inter-individual variation rather than pregnancy progression, consistent with our use of a broad-scope proteomics panel not specifically designed to capture pregnancy- or labour-related signals. Full details of the exploratory analyses are provided in **Supplementary Note 2**.

To identify proteins with consistent temporal trends preceding delivery while accounting for inter-individual variability, we fitted one generalized additive mixed model (GAMM) per protein, treating days before delivery as the fixed effect and individual as a random effect (**Supplementary Note 3**). This approach identified 28 proteins with strong temporal trends in the exploratory group (conditional R2 ≥ 0.7 that replicated in the confirmatory group under the same criterion. The trajectories spanned a range of temporal patterns, with some proteins displaying largely linear trends throughout the investigated period, while others showed non-linear dynamics concentrated in specific time windows closer to delivery.

Strikingly, the individual-level trajectories of the former (IL1RL1, ANGPT2, and AFP) showed near-universal agreement across women, with protein levels in almost all individuals changing in the same direction with similar slopes, indicating that signals from these proteins reflect a highly consistent biological programme. In contrast, proteins from the latter group (e.g., MMP8, MMP9, IL6 and OSM) showed a sharp increase in abundance only in the four days before delivery, consistent with a coordinated terminal inflammatory effector programme, but with considerably greater inter-individual variability in the timing and magnitude of this rise compared to the other signals. More details on the biological role of these four proteins are presented in **Supplementary Note 3**.

### Machine learning predicts time-to-delivery from plasma proteomics

Gradient-boosted decision tree models (XGBoost) trained on a dataset including all 5,416 proteins and gestational age achieved a mean leave-one-subject-out (LOSO) root mean square error (RMSE) of 2.60 ± 1.75 days and mean absolute error (MAE) of 2.18 ± 1.60 days in the exploratory group. A proteomics-only model (RMSE 3.05 ± 1.05) outperformed random forest, lasso regression, elastic net, and a baseline assuming delivery at the expected date of 40+0 weeks (**Supplementary Note 4**), indicating that protein signatures captured aspects of the timing of labour onset beyond gestational age alone. The model including gestational age generalized to the confirmatory group. SHAP analysis – which decomposes each model prediction into additive protein-level contributions – identified a consistent set of protein predictors across both groups (**Supplementary Note 5**) with positive SHAP contributions corresponding to increasing protein levels closer to delivery (e.g. IL1RL1, ACTA2, LMOD1) and negative contributions to declining protein levels (e.g. AFP, ANGPT2), confirming directional concordance between machine learning signals and longitudinal trajectories.

Predictive accuracy was highest in the final days before delivery, when several proteins underwent dramatic changes in level (e.g. ACTA2, LMOD1, proteins discussed in **Supplementary Note 3**). In this time window, good model performance likely reflects detection of a labour initiation sequence already underway at a subclinical level, rather than prediction of future onset. Early and sustained changes present weeks before delivery, such as those observed for IL1RL1, ANGPT2 and AFP, may better support advance prediction of labour onset.

### Five proteins spanning feto-placental, vascular, immune and contractile biology show consistent, cross-method support for key roles in labour onset

Five proteins showed consistent, cross-method support for key roles in initiating labour, appearing as top contributors in both the XGBoost SHAP analysis and the GAMM longitudinal modeling in both the exploratory and confirmatory groups: AFP, ACTA2, ANGPT2, IL1RL1, and LMOD1 (**Table 2**). These proteins span a range of biological functions and compartments – feto-placental, vascular, immunological, and contractile – and their convergence across two independent analytical frameworks provides strong evidence that they capture genuine biological signals of imminent labour rather than model-specific artifacts.

**Table 2.** Proteins with consistent cross-method and cross-group support for key roles in initiating labour. For each protein we report gene symbol, UniProt AC, XGBoost SHAP importance in the exploratory group and the corresponding GAMM conditional R^2^ (GAMM cR^2^). The same metrics are shown for the confirmatory group. SHAP importance reflects the contribution of each protein to XGBoost prediction of days-to-delivery, with higher values indicating stronger association with the outcome. GAMM cR^2^ is a measure of longitudinal trajectory consistency, reflecting how regularly and consistently a protein changes over time across participants, thus a high GAMM cR² indicates a protein with a reproducible, low-noise temporal trajectory.

| Gene name | UniProt AC | Exploratory group |  | Confirmatory group |  |
| --- | --- | --- | --- | --- | --- |
| | | SHAP | GAMM $cR^2$ | SHAP | GAMM $cR^2$ |
| AFP | P02771 | 0.74 | 0.76 | 0.63 | 0.82 |
| ACTA2 | P62736 | 0.60 | 0.95 | 0.66 | 0.88 |
| ANGPT2 | O15123 | 0.70 | 0.70 | 0.70 | 0.87 |
| IL1RL1 | Q01638 | 0.23 | 0.77 | 0.23 | 0.84 |
| LMOD1 | P29536 | 0.59 | 0.97 | 0.59 | 0.92 |

Levels of alpha-fetoprotein (AFP), a glycoprotein produced by the fetal yolk sac and liver, decline progressively toward term in maternal circulation^11^, reflecting a gradual withdrawal of this fetal protein^12^ as the placenta ages^13^. We interpret AFP not as a mechanistic participant in labour initiation, but as an indicator that the feto-placental unit has reached the maturation threshold that permits, without directly triggering, labour onset.

Angiopoietin-2 (ANGPT2) acts as a natural antagonist of the Tie2 receptor, promoting endothelial destabilization and vessel remodeling^14^, and is expressed in the human placenta and trophoblast throughout gestation^15,16^ and up to labour^17^. This protein shows a monotonic decline in the weeks preceding delivery (**Figure 2**), which can be explained by two complementary mechanisms. First, ANGPT2 expression is regulated by progesterone during pregnancy^18^, and as a downstream consequence of the hormonal transition preceding labour, progesterone withdrawal near term reduces ANGPT2. Second, the transition from active vascular remodelling to pre-contractile vascular stability reflects uterine preparation for the hemodynamic demands of labour^19^.

**Figure 2.**
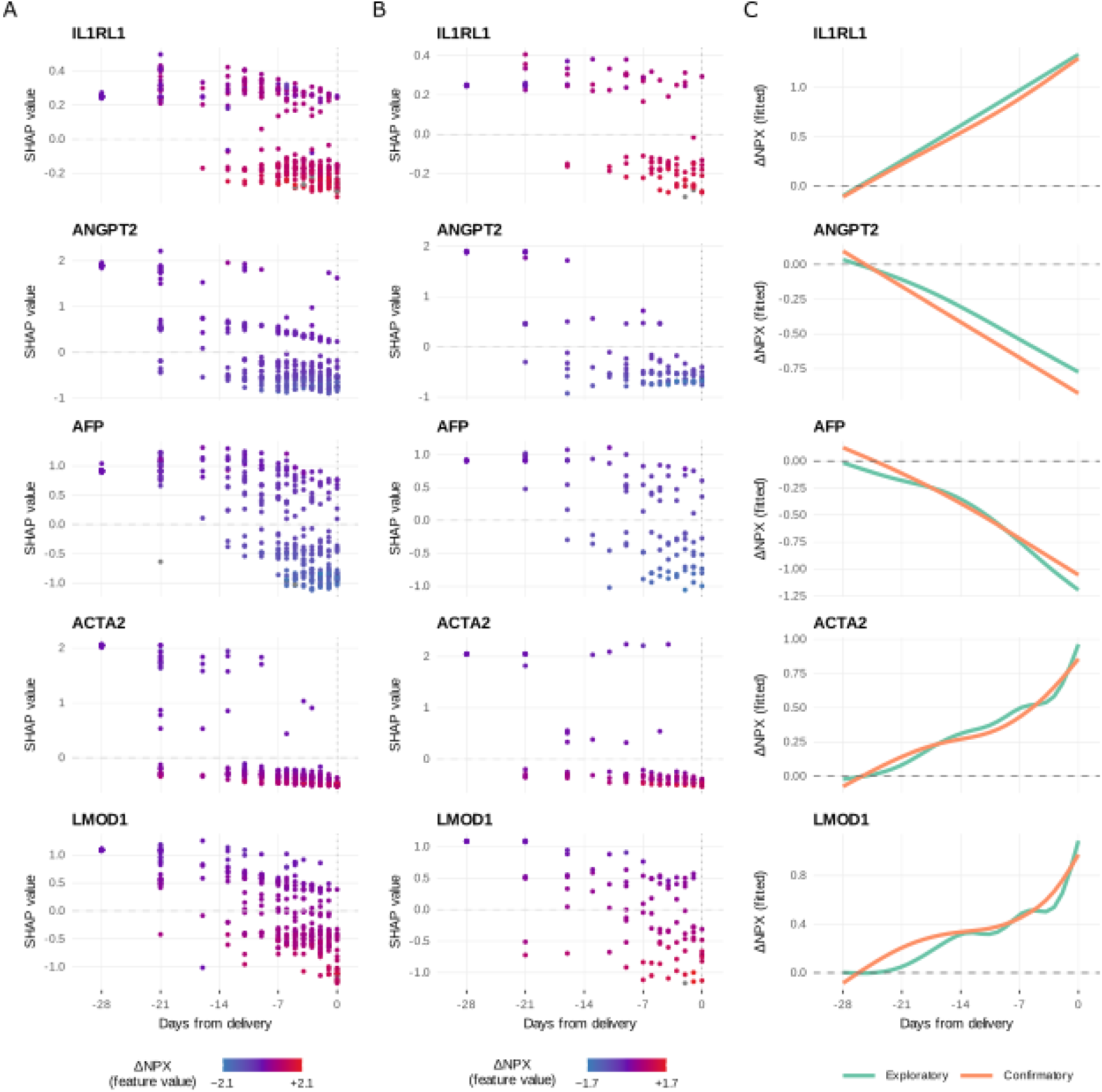
SHAP dependency plots of plasma protein dynamics preceding labour onset in the (A) exploratory and (B) confirmatory groups. Each panel shows individual plasma samples plotted by days from delivery (x axis) and SHAP value, i.e. the contribution of that protein in that sample to the XGBoost prediction of days from delivery (y axis), with point colour indicating the feature value (ΔNPX, change in protein expression from each woman’s individual baseline). Red denotes a high, positive ΔNPX (rising protein level) and blue denotes a low, negative ΔNPX (falling protein level). The dashed horizontal line marks SHAP = 0 and the dotted vertical line marks the day of delivery. Panels are shown for the 5 key proteins identified by both XGBoost models and GAMMs (IL1RL1, ANGPT2, AFP, ACTA2, LMOD1). **(C) GAMM fitted smooth trajectories for each protein across the antepartum window, estimated from the full proteomics dataset using one model per protein and random effects for individual women.** Mint green lines show the exploratory group and orange lines show the confirmatory group. The y axis shows the fitted ΔNPX from individual baseline and the x axis corresponds to days from delivery. The dashed horizontal line marks ΔNPX = 0. All five proteins show consistent directional trajectories across both groups, with ANGPT2 and AFP declining and IL1RL1, ACTA2 and LMOD1 rising in the weeks preceding delivery.

IL1RL1, encoding both soluble ST2 (sST2) and membrane-bound ST2L via alternative splicing, showed the earliest and most consistent rise across all women in our study (**Figure 2**). sST2 is the competitive inhibitor for IL-33, a nuclear alarmin constitutively expressed in cervical and uterine epithelial cells^20^. The ratio of sST2 to membrane bound ST2L determines net IL-33 availability^21,22^. Prior work has established that the IL-33/ST2 axis is biologically active in the reproductive compartment and functionally relevant to labour timing: (i) ST2 and IL-33 are expressed in human placenta, decidua, and trophoblast^23^, (ii) sST2 is measurable in maternal plasma across gestation and shows longitudinal variation associated with pregnancy complications^24^, and (iii) parturition pathways in mice have been shown to depend upon innate type 2 immunity^25,26^, which is activated by IL-33 through its action on group 2 innate lymphoid cells, mast cells, and Th2 cells^27,28^, providing direct mechanistic support that the IL-33/ST2 signaling axis is relevant to labour timing.

The near complete absence of free IL-33 in peripheral plasma (490/491 samples were below the limit of detection) alongside rising sST2 levels is consistent with progressive IL-33/ST2 pathway activation^20^, in which co-induced sST2 dynamically regulates free IL-33 availability as the pathway escalates. The concentrations required for autocrine or paracrine ST2L activation should be substantially lower than those required for peripheral plasma accumulation^29^. This, in conjunction with the short extracellular half-life of IL-33 (rapid oxidation, proteolytic cleavage, sST2 binding),^30^ makes systemic accumulation unlikely even during active local signaling. In cardiovascular settings, where cardiac fibroblasts under mechanical strain co-induce both IL-33 and sST2^31^, sST2 has been validated as a biomarker of IL-33 pathway activation in heart failure, where free IL-33 remains undetectable even during active signaling, making sST2 the accessible readout of pathway activity^22,31^. Full mechanistic and interpretive considerations for the IL-33/ST2 findings, including nuclear retention of IL-33, and alternative hepatic acute-phase response interpretations for rising sST2, are addressed in **Supplementary Note 6**.

Smooth muscle α-actin (ACTA2) and leiomodin-1 (LMOD1) showed a consistent late rise in both groups, positioning these proteins as downstream effectors of IL-33/ST2 alarmin signaling and myometrial contractile priming. ACTA2 is a canonical marker of smooth muscle cell differentiation and activation, with confirmed expression in the myometrium throughout pregnancy^32^. LMOD1 is an actin filament nucleator, expressed preferentially in differentiated smooth muscle cells^33,34,35^, and the Human Protein Atlas^36^ confirms its expression in smooth muscle, endometrium, cervix, and placenta. To our knowledge, circulating LMOD1 has not previously been reported as a plasma biomarker in the context of pregnancy or labour. Notably it achieved the highest GAMM conditional R² in both the exploratory (0.97) and confirmatory (0.92) groups, indicating exceptional longitudinal predictability. We propose that rising plasma LMOD1 reflects progressive assembly of the contractile apparatus in smooth muscle tissue, potentially including the myometrium and cervical smooth muscle, reaching peripheral plasma through extracellular vesicle release from activated smooth muscle cells, an established route by which intracellular proteins reach systemic circulation during tissue activation^37,38,39^.

Network and enrichment analysis using STRING^40^ confirmed that the interaction neighbourhoods of all five proteins were consistent with their proposed biological roles (**Supplementary Note 7**).

### The five proteins change in a sequential order consistent with a biological cascade

To directly visualize the temporal dynamics of the five key proteins, we plotted their SHAP dependency scores against time, coloured by feature value (ΔNPX) alongside their fitted smooth GAMM trajectories (**Figure 2**). Since XGBoost captures feature interactions, two samples with identical ΔNPX could carry different SHAP values depending on the time between sample collection and delivery. A protein is contributing strongly to predictions at a given timepoint when protein levels are changing across women (colour gradient along the y-axis at that timepoint) and those level differences are driving correspondingly different predictions (vertical spread in SHAP values at that timepoint). When colour gradient and SHAP spread co-occur, this indicates that the protein is both changing in level and maximally informative to the model at that moment. This temporal structure differed across proteins. For IL1RL1, the colour gradient, despite not exhibiting big fluctuations, was present and consistent across the last five weeks of pregnancy, reflecting its early and sustained rise throughout that period. For ACTA2, the strongest colour gradient emerged ∼21 days before delivery and attenuated in the final week, indicating that its predictive contribution peaks during 21–7 days before delivery. For AFP and ANGPT2, the gradient intensified progressively closer to delivery, consistent with a decline that accelerates as term approaches. LMOD1 showed a concentrated band of high-SHAP, high ΔNPX points in the final days before delivery. All five proteins showed directionally consistent changes across the sampling window, with ACTA2 and IL1RL1 rising monotonically and AFP and ANGPT2 falling monotonically as delivery approached. LMOD1 showed a marked increase in abundance shortly before labour onset. The concordance between SHAP contributions and GAMM-fitted trajectories confirms that the XGBoost model recovered the same biological signal identified by the trajectory modelling, and that these signals generalized consistently to the confirmatory group.

Directional consistency analysis confirmed that the expected direction of change was observed in at least 29 of the 30 women in the exploratory group and all 10 women in the confirmatory group for all five proteins, with all proportions significantly exceeding the permutation-derived null distribution (p<0.001 for all proteins, **Figure 3A**). We also evaluated our findings in three publicly available independent datasets that were as similar to ours as possible. Although no existing public dataset matched the combination of high-dimensional proteomics, dense third-trimester longitudinal sampling, and delivery anchored temporal resolution that characterized our population, both IL1RL1 and ANGPT2 trajectories were independently replicated across all populations and proteomic platforms^6–8^ (IL1RL1 87% directional consistency, p<0.001; ANGPT2 79%, p<0.001 in Tarca et al. 2022^6^; **Supplementary Note 8**).

**Figure 3.**
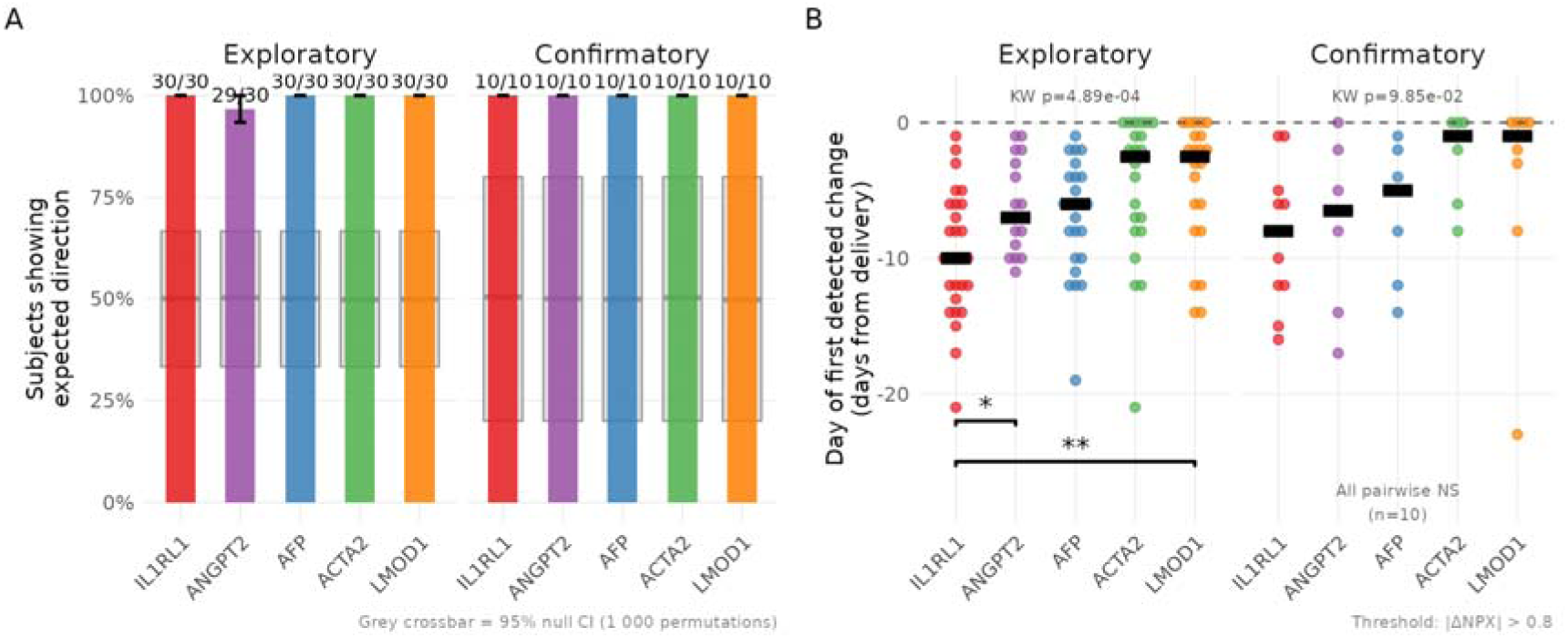
**(A) Proportion of women showing a change in the level of each of the five key proteins in the expected direction** (rise for IL1RL1, ACTA2, LMOD1; fall for AFP, ANGPT2), **computed from first to last available timepoint per woman**. Fractions above the bars indicate the number of women whose protein level changed in the expected direction over the total number. Grey crossbars represent the 95% null confidence interval derived from 1,000 permutations in which NPX values were randomly shuffled within each woman-protein combination, preserving sample size and timepoint structure while destroying the temporal signal. Permutation-derived p-values for all tests were ≤ 0.001. **(B) Day of first detected change for each protein, defined as the earliest timepoint at which** Δ**NPX exceeded ±0.8 NPX units in the expected direction.** Each dot represents one woman; horizontal black bars indicate the median. Proteins are ordered along the x axis according to their hypothesized cascade position. The Kruskal-Wallis p-value testing whether onset timing differed across proteins is shown in each panel. In the exploratory group, pairwise Wilcoxon tests with Benjamini-Hochberg correction indicated that IL1RL1 levels began to change significantly earlier than levels of both ANGPT2 (p=0.047) and LMOD1 (p=0.004); no other pairs differed significantly in timing of change onset.

Crucially, levels of the five proteins did not change simultaneously, but in a statistically significant sequential pattern (**Figure 3B**). The robustness of the overall ordering signal to the choice of threshold was assessed in a sensitivity analysis using protein-relative thresholds (**Supplementary Note 9**). IL1RL1 showed the earliest median onset of change, preceding ANGPT2 and AFP, which in turn preceded the contractile proteins ACTA2 and LMOD1. This ordering was statistically significant by Kruskal-Wallis test in both the exploratory and confirmatory groups. Pairwise Wilcoxon tests with Benjamini-Hochberg correction confirmed that onset of changes in IL1RL1 levels occurred significantly earlier than in ANGPT2 and LMOD1 in the exploratory group. Pairwise comparisons in the confirmatory group were all non-significant, likely reflecting the smaller sample size.

Integrating the temporal order of changing protein levels with the biological identity of each protein, we propose a two-phase cascade model of spontaneous labour onset at term (**Figure 4**). In an early priming phase (approximately 1–4 weeks before labour), feto-placental aging drives declining AFP levels and functional progesterone withdrawal reduces ANGPT2 levels as the uterine vasculature transitions towards pre-contractile stability^18^. IL-33 released from stressed reproductive tissues co-induces rising sST2 (IL1RL1) – reflecting escalating IL-33/ST2 pathway activity – while the sST2/ST2L balance modulates the rate of downstream membrane-bound signaling. In a subsequent execution phase (final days before labour) this balance shifts – whether through declining sST2 relative to ST2L, rising IL-33 release, or both – and free IL-33 signals through membrane-bound ST2L on myometrial and immune cells; ACTA2 and LMOD1 rise as the contractile apparatus assembles. The late phase is further amplified by a sharp rise in IL-6 and OSM (gp130 cytokine axis) and MMP8 and MMP9, driving prostaglandin synthesis and cervical remodelling in the final days before delivery. Consistent with this model, IL-33 has been reported in the nuclei of smooth muscle cells in the cardiac vasculature, where ST2 is co-expressed^41^, raising the possibility that IL-33 released from stressed smooth muscle cells in the uterus could signal in an autocrine or paracrine fashion through membrane-bound ST2 on adjacent myometrial cells, driving cytoskeletal reorganization and LMOD1 upregulation as part of the contractile priming program. Taken together, all these signals constitute an integrated inflammatory effector programme in which the alarmin cascade, initiated weeks before labour, converges on a final common pathway of cervical remodelling and myometrial activation.

**Figure 4.**
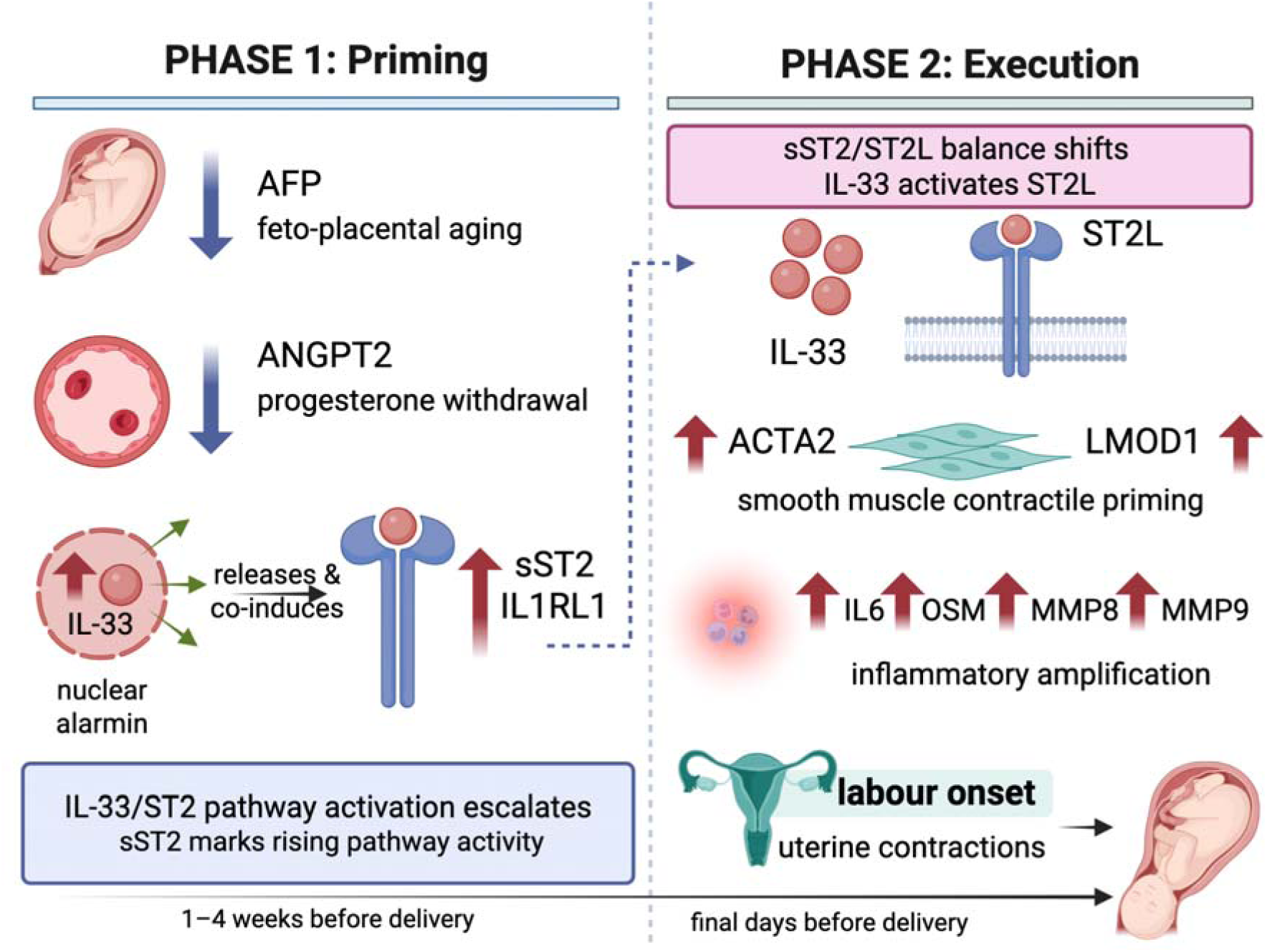
Proposed two-phase biological cascade preceding spontaneous labour onset at term. In Phase 1 (priming; 1–4 weeks before delivery), feto-placental aging drives a progressive decline in AFP levels, while functional progesterone withdrawal reduces ANGPT2 levels, reflecting the transition of the uterine vasculature toward pre-contractile stability. Concurrently, IL-33 progressively co-induces rising soluble ST2 (encoded by IL1RL1); rising plasma sST2 reflects escalating IL-33/ST2 pathway activity. In Phase 2 (execution, final days before delivery), the sST2/ST2L balance shifts in favour of membrane signaling, and free IL-33 drives paracrine signaling through ST2L. This triggers upregulation of the smooth muscle contractile proteins ACTA2 and LMOD1, reflecting the assembly of the myometrial contractile apparatus. A terminal inflammatory amplification programme – marked by sharp rises in IL-6, OSM, MMP8 and MMP9 – drives prostaglandin synthesis and cervical remodelling, culminating in uterine contractions and labour onset. Arrows denote direction of change in protein levels (up arrow: rising, down arrow: falling) as delivery approaches. A dashed arrow indicates the proposed mechanistic transition between the two phases. AFP: alpha-fetoprotein, ANGPT2: angiopoietin-2, IL-33: interleukin 33, sST2/IL1RL1: Soluble Suppression of Tumorigenicity 2/Interleukin-1 receptor-like 1, ST2L: membrane bound ST2, ACTA2: smooth muscle α-actin, LMOD1: leiomodin 1, IL-6: Interleukin 6, OSM: oncostatin M, MMP: matrix metalloprotease.

### Mendelian randomization analysis of overlapping proteins and gestational duration

To assess whether the longitudinal protein changes observed in our population reflect causal biological processes rather than correlated markers of gestational age, we performed two-sample Mendelian randomisation^42^ (MR) for each of the five overlapping proteins against three gestational outcomes from the Early Growth Genetics Consortium^43^. Of the five proteins only IL1RL1 yielded statistically significant results: genetically higher sST2 levels were associated with shorter gestational duration (β=−0.51 days per SD, 95% CI: −0.90 to −0.12, p=0.010, FDR=0.04), with no evidence of heterogeneity across the six independent genetic variants (Cochran’s Q=8.18, df=5, p=0.147) or of directional pleiotropy (MR-Egger intercept=0.016, p=0.903). To confirm that the genetic variants used are acting specifically through IL1RL1 rather than through unrelated biological pathways we queried all variants against a database of genome-wide associations (PheWAS). The only non-proteomic associations identified were with eosinophil counts, asthma, atopic dermatitis and inflammatory bowel disease, all of which are established downstream consequences of IL-33/ST2 signaling, consistent with the variants acting specifically on this pathway. ANGPT2 showed no causal effect on gestational timing across any outcome (β=+0.021 days, p=0.94). No significant MR result was identified for AFP, ACTA2 or LMOD1. Because genetic variants are fixed at conception and cannot be caused by the pregnancy itself, these results suggest that higher IL1RL1 levels are a determinant of labour timing rather than a downstream marker of it. Full results for all proteins across all outcomes, sensitivity analyses, and instrument validation are provided in **Supplementary Note 10**.

## Discussion

We present a longitudinal plasma proteomics study of 40 women with spontaneous labour at term, using a two-stage analytical pipeline combining GAMM-based trajectory modeling, machine learning prediction, and SHAP-based interpretability to identify proteins associated with labour timing. We identified a reproducible set of five proteins (AFP, ACTA2, ANGPT2, IL1RL1, LMOD1) with consistent cross-method and cross-group support. The temporal ordering of these proteins is consistent with a sequential biological cascade preceding labour onset that we propose is initiated by the IL-33/ST2 alarmin axis.

The initiation of spontaneous human labour has long been recognized as an inflammatory process, but the upstream triggers that activate this inflammation in the absence of infection remain incompletely understood^1^. Sterile intra-amniotic inflammation has been documented in a significant proportion of women presenting with preterm labour with intact membranes^44^, and a similar sterile inflammatory program appears to characterize the transition to term labour^1^. The conceptual framework for how sterile inflammation can be initiated and sustained is provided by the damage-associated molecular pattern (DAMP) model, in which endogenous molecules released from stressed, damaged, or senescent cells activate innate immune receptors in the absence of pathogen-associated signals^45^. Multiple DAMPs have been detected in reproductive compartments at the time of parturition, including HMGB1^46,47^, S100A12^48^ and serum amyloid A^49^, and fetal membrane senescence^50,51^ has been proposed as a fundamental timing mechanism for parturition^52^. Our data extend this framework by identifying IL-33, a nuclear alarmin expressed in cervical and uterine epithelial cells^20^, as a candidate upstream initiating signal, with plasma proteomic evidence of rising IL1RL1/sST2 reflecting escalating IL-33/ST2 pathway activity in the weeks preceding labour.

The cascade model integrates converging evidence from longitudinal proteomics, machine learning feature importance, GAMM trajectory modelling, cross-population replication, and human genetics into a coherent biological narrative of spontaneous labour onset at term. The causal role of the IL33/ST2 axis is grounded on an accumulating body of literature^23,25,26,28,53^ and is directly supported by Mendelian randomisation analyses showing that genetically higher sST2 levels shorten gestation, i.e., by human genetic evidence consistent with a role for the IL-33/ST2 axis in labour timing beyond mere correlation. The mechanistic links between individual proteins remain to be confirmed by direct experimental tests, and the proposed ordering reflects the most parsimonious interpretation of temporal and genetic evidence rather than established causality at each step. The model makes specific testable predictions – most importantly, that IL-33 levels in decidua or amniotic fluid should show a temporally consistent rise in the weeks before term labour, that experimental blockade of sST2 should accelerate or potentiate labour onset in animal models, and that pharmacological or genetic attenuation of the gp130 cytokine axis or MMP activity in the final days of gestation should delay cervical ripening without affecting the earlier priming phase.

## Conclusions

We have identified a reproducible five-protein signature of labour proximity in maternal plasma, characterized by an early rise in IL1RL1, a concurrent fall in the vascular remodelling factor ANGPT2 and fetal protein AFP, and a late rise in the smooth muscle contractile proteins ACTA2 and LMOD1. The temporal ordering of these signals is consistent with a sequential biological cascade in which feto-placental aging and vascular maturation create the conditions for IL-33/ST2 alarmin axis activation, reflected in rising plasma sST2, until the sST2/ST2L balance shifts toward membrane-bound signaling, at which point free IL-33 drives myometrial priming and labour execution.

## Methods

### Biological Signals in Pregnancy cohort

The Biological Signals in Pregnancy (BSG) cohort was established in Copenhagen, Denmark, between 2014 and 2019^54,55^. Participants were enrolled via referrals from family doctors and advertisements. At enrolment, women were screened to confirm good general health, absence of chronic disease, and no current medication use. Maternal age at delivery ranged from 23 to 36 years. Non-fasting blood samples were collected weekly throughout pregnancy and once postpartum (two 9-mL EDTA tubes and one PAXgene RNA tube per draw). a subset of the cohort also provided daily samples after 38 completed weeks of gestation.

The present study used a sub-cohort of women from BSG who had reached 36+0 gestational weeks without complications and were enrolled in a dense sampling protocol, with blood samples collected at 36, 37, and 38 weeks, and then daily from 39+0 weeks until delivery, with one additional sample collected in active labour and one postpartum. From this sub-cohort, women were selected for the present analysis if they had a known delivery date and underwent spontaneous labour onset, with or without augmentation after spontaneous onset. Women who underwent acute caesarean section without spontaneous labour onset or for whom delivery date could not be confirmed were excluded, resulting in a final study population of 40 women.

To ensure a balanced division of participants for model development and validation, we stratified the population into exploratory (n = 30) and confirmatory (n = 10) groups using k-means clustering. Stratification was based on maternal age at delivery, gestational age (GA) at delivery, and parity. These variables were scaled and used as input to a 3-cluster k-means model, preserving diversity across key maternal characteristics. Within each cluster, 75% of individuals were randomly sampled to form the exploratory group, with the remaining participants allocated to the confirmatory group. This design ensured representative coverage of relevant clinical covariates in both groups (Table 1).

### Ethical approval and consent to participate

The study was approved by the Scientific Ethics Committee for the Capital Region of Denmark (H-3-2014-004) and Statens Serum Institut’s Department of Data Protection and Information Security and reported to the Danish Data Protection Agency. All participants provided written informed consent.

### Proteomic Profiling and Preprocessing

Samples were randomized across 96-well plates, with the constraint that all longitudinal samples from the same woman were assigned to the same plate. Positions were reserved on each plate for external quality controls. Protein expression profiling was performed using the Olink Explore HT platform, which employs proximity extension assay technology to simultaneously measure 5,416 protein targets with high specificity and sensitivity^56^. The output consisted of NPX values on a log_2_ scale, which represent relative quantification across samples. Olink’s standardized preprocessing pipeline, including normalization and batch correction, was applied.

### Study Design and Data Processing

Of the 2,671,040 protein-sample combinations in the full NPX dataset, 85.4% (n=2,282,158) were retained for analysis, comprising antepartum samples from women with known delivery dates and spontaneous labour onset, with valid UniProt identifiers and non-zero NPX values. Following this step, NPX values were normalized within each individual by subtracting the NPX value of their earliest antepartum sample:

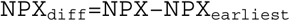

This transformation allowed the modeling of protein trajectories relative to an individual-specific baseline.

To address variability in sampling schedules across participants, we discretized all sample timepoints into a common temporal framework based on days before delivery. This approach enabled alignment of protein expression measurements across individuals, facilitating robust trajectory modeling. We used a tiered binning strategy that captured distinct windows of late gestation. Specifically, in the early phase of sampling (≥15 days before delivery), timepoints were coarsely grouped to accommodate sparser sampling. Specifically, all samples collected 25 or more days before delivery were assigned to a bin labelled “–28”. Samples taken between 24 and 18 days before delivery were grouped under “–21”, while those between 17 and 15 days were grouped as “–16”. During the mid phase (–14 to –7 days), sampling density increased. Here, timepoints were binned into 2-day intervals by rounding each day upward to the nearest odd number. For instance, both –10 and –9 were grouped into –9, ensuring consistent alignment across subjects while preserving some temporal resolution. In the late phase (–6 to 0 days), where changes in protein expression were expected to be more dynamic and sampling was most dense, each day was treated as a separate bin. This fine-grained resolution allowed for detailed modeling of the immediate pre-labour period.

When multiple samples for the same protein fell within the same bin for an individual, a single representative measurement was selected using a biologically informed prioritization scheme. Preference was given to samples whose timing exactly matched the bin midpoint, and among ties, to samples collected on days with higher overall sampling density. This strategy ensured that selected data points were both temporally representative and well-supported. The final grouping provided a harmonized time axis for modeling protein trajectories across the group and only 3 samples were excluded by the process.

### Modeling Protein Expression Over Time

To characterize protein expression dynamics over time, we fit generalized additive mixed models (GAMMs)^57^ independently for each protein using the mgcv package v. 1.9-3 in R 4.5.0. GAMM with random intercepts were defined as:

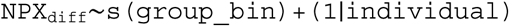

Models were fitted using mgcv::gamm(). Fixed effects were estimated by using a smooth function of the time point groups, using a thin plate regression spline for the time point groups’ effect. Random effects were specified separately to account for individual-level variability (i.e., between-woman variability). The smoothness of the time function was determined via restricted maximum likelihood (REML). The maximum number of spline basis functions was adaptively constrained to 8 or fewer, based on the number of unique bin levels for each protein. For each model, we computed marginal and conditional R^2^. Finally, we performed 5-fold cross validation to assess the generalizability of our models.

We first ranked the GAMM models by conditional R^2^, which includes both fixed and random effects. We further considered only models with conditional R^2^≥0.7, to ensure that only good quality models would be retained moving forward. We have performed the same analysis both for the exploratory group and retained only those models with conditional R^2^≥0.7 in both groups. More details on this analysis are available in **Supplementary Note 3**.

### Machine learning for labour timing

The dataset was transformed into a sparse matrix of dimensions n x p, where rows corresponded to unique samples and columns to proteins, to allow training of gradient boosting models. The response variable was defined as the “absolute number of days from delivery at the time of sampling”. Because multiple samples were available per woman, we adopted grouped cross-validation (CV) to prevent information leakage. Two levels of CV were employed. In the first stage, a 5-fold grouped CV was used for hyperparameter tuning where women were randomly partitioned into five folds, with all samples from the same woman assigned to the same fold, ensuring that samples from no woman appeared in both training and validation sets for any split. This setup was used within Bayesian optimization^58^ to efficiently explore hyperparameters. The outcome was absolute number of days to delivery at sampling. In the second stage, the tuned hyperparameters were used in a leave-one-subject-out (LOSO) cross-validation for final model evaluation, in which all samples from one woman were held out in turn while the model was trained on the remaining women. This two-stage design prevents data leakage at both the hyperparameter selection and performance estimation steps.

Gradient boosting models were trained using XGBoost^59^ with the squared error loss. The following hyperparameters were optimized: learning rate (η), maximum tree depth, subsample ratio, column subsampling ratio, minimum child weight, and L1/L2 regularization terms (α, λ). Bayesian optimization was performed with 10 random initialization points and 40 acquisition rounds using an upper confidence bound acquisition function (κ = 2.576). Each trial was evaluated with up to 2,000 boosting rounds and an early stopping criterion of 100 rounds. The tuned hyperparameters were then used in the second stage of cross-validation (LOSO), in which all samples from one woman were held out in turn, the model was trained on the remaining women and was evaluated on the held-out woman. Root mean square error (RMSE) and mean average error (MAE) were computed per fold and averaged across folds.

For the final model, we appended gestational age at sampling as an additional predictor to the proteomic feature set and refit the model under the same grouped CV and Bayesian optimization protocol on the exploratory group. Gestational age (GA, in days) was computed per sample as:

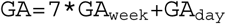

standardized within each training split to avoid leakage.

To contextualize performance, XGBoost was benchmarked against three alternative predictors: (a) Random Forest (1000 trees, mtry = √p, min node size = 5), (b) Lasso regression (α = 1) with penalty parameter chosen by internal 5-fold CV, and (c) Elastic Net regression (α = 0.5) with penalty parameter chosen similarly.vMoreover, we compared the best model against a deterministic baseline of *40 weeks+0 days*, which assumes all women deliver exactly 280 days after their last menstrual period. Expected date of delivery was computed based on the woman’s first trimester ultrasound scan.

All models were compared on RMSE and MAE across LOSO folds. For the deterministic baseline, RMSE was reported without variability estimates and predicted days-from-delivery were computed as:

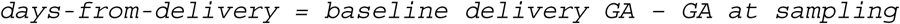

RMSEs were calculated across all available samples. The final model was trained with up to 300 boosting rounds, using an early stopping criterion of 20 rounds to prevent overfitting.

To interpret the model, we applied SHapley Additive exPlanations (SHAP)^60^, which decomposes each prediction into additive feature contributions. SHAP values were computed for every sample in the dataset. These values quantify how much each protein increased or decreased the predicted time to delivery relative to the model’s baseline prediction (bias term).

Feature importance was assessed by calculating the mean absolute SHAP value per protein across all samples (global importance). This identifies proteins that consistently exert strong influence on predictions. We visualized the global importance with a SHAP beeswarm plot, displaying both the magnitude and direction of effects for the top-20 features, colored by feature value (ΔNPX).

### Evaluation on the confirmatory group

To assess whether the proteomic signature of labour proximity identified in the exploratory group generalizes to held-out data, we evaluated the final XGBoost models trained on the full exploratory proteomics dataset with and without GA, using the best hyperparameters identified via Bayesian optimization on the held-out confirmatory test group (n = 10 participants). The dataset from the confirmatory group was processed identically to the exploratory group, with the same protein filtering and normalization steps and construction of individual-level NPX differences. All feature columns were aligned to the exploratory training space to ensure consistent dimensionality, and no re-training or re-tuning was performed on the confirmatory data. Model performance was quantified using RMSE and MAE between predicted and observed absolute days-from-delivery. Feature attributions were computed for all confirmatory samples using SHAP, following the same procedure applied to the exploratory data, to evaluate reproducibility of global feature importance rankings.

### Directionality consistency and temporal ordering analysis

For each protein and each woman, the observed direction of change was defined by comparing the ΔNPX value at the last available timepoint to the first available timepoint within the antepartum sampling window. A woman was classified as showing change in the expected direction if the observed change matched the hypothesized direction for that protein (IL1RL1, ACTA2, LMOD1: upward; AFP, ANGPT2: downward). The proportion of women with change in the expected direction was computed per protein per group, with a standard error derived from the binomial variance formula. To assess whether the observed proportions exceeded chance expectation, a permutation null distribution was generated by randomly shuffling ΔNPX values within each woman-protein-group set across 1000 permutations, preserving the temporal structure but breaking the directional signal. The 2.5^th^ and 97.5^th^ percentiles of the permuted proportions were used as the 95% null confidence interval, and a permutation p-value was computed as the proportion of permuted values equal to or exceeding the observed proportion.

The day of first detectable protein change for each protein in each woman was defined as the earliest timepoint at which the absolute ΔNPX exceeded a threshold of 0.8 NPX units in the expected direction. This threshold was chosen to distinguish a meaningful change from baseline noise, thus women whose protein levels changed in the expected direction but never exceeded this threshold – reflecting a smaller absolute excursion rather than absence of change – could not be assigned an onset day of change and were therefore excluded from the onset timing analysis. The median onset day of change was computed per protein per group and displayed as a crossbar. To test whether the five proteins showed different onset timing, a Kruskal-Wallis test was applied separately within each group. Pairwise comparisons between proteins were performed using Wilcoxon rank-sum tests with Benjamini-Hochberg correction. Significant pairwise comparisons in the exploratory group are annotated on **Figure 3**. No pairwise tests were performed in the confirmatory group due to the small sample size (n=10).

### Mendelian Randomization Analysis

To assess causality, we performed two-sample MR^42,61^ using the TwoSampleMR R package. Genetic instruments were obtained as cis-pQTLs from Loya et al., 2025^62^, a genome-wide association study (GWAS) of 2,923 plasma proteins measured by Olink Explore 3072 PEA in 47,745 UK Βiobank participants of European ancestry ^63^. Each protein variant within 1 Mb of the gene transcription start site (GRCh37) reaching genome-wide significance (p<5×10^-8^) was selected and clumped for independence (r^2^<0.001, 10 Mb window, EUR linkage disequilibrium reference^61^). Instruments with F-statistics below 10 were excluded to guard against weak instrument bias^64^. The primary estimator was inverse variance weighted (IVW) meta-analysis for proteins with multiple instruments (genetic variants), and the Wald ratio for proteins with a single instrument. Sensitivity analyses for proteins with three or more instruments included the weighted median estimator and MR-Egger regression^65^. Heterogeneity across instruments was assessed using Cochran’s Q statistic and the MR-Egger intercept test was used to assess directional pleiotropy. The Steiger filtering test confirmed the causal direction ran from protein level to gestational duration rather than the reverse. FDR correction was applied separately within continuous outcomes and binary outcomes using the Benjamini-Hochberg procedure. Three gestational outcomes from the

Early Growth Genetics Consortium^43^ were tested: gestational duration (days, n=151,987), post-term delivery (≥42 weeks, n=131,279; cases=15,972), preterm delivery (<37 weeks, n=233,290; cases=15,419). All instrument SNPs were queried against the IEU OpenGWAS phenotype-genotype map (p<10^-5^) to identify pleiotropic associations that could violate the exclusion restriction.

All analyses were performed in R (v4.3). The following packages were used: umap, colorspace, xgboost, ranger, glmnet, rBayesianOptimization, Matrix, dplyr, tidyr, doParallel, lme4, lmerTest, foreach, ggplot2, readr, ggbeeswarm, forcats, RColorBrewer, purrr, ggsignif, scales, tibble, OlinkAnalyze, patchwork, lubridate, stringr, tidyverse, survival, RCy3, GEOquery, TwoSampleMR, ieugwasr, openxlsx, gt, MuMIn, rlang, eulerr.

## Acknowledgements

We are immensely grateful to the dedicated women who participated in the study, generously donating blood samples during their pregnancy. We would also like to thank everyone involved in data collection and biological material handling.

This work was supported by Independent Research Fund Denmark (9039-00417B). KN and HAB were supported by the Novo Nordisk Foundation (NNF19OC0054286).

## Author contributions

KN, MT, HAB and MM conceptualized and designed the study with input from the remaining authors. NMS, M-LHR and MM organized and contributed to the collection of pregnancy samples. KN and MT analyzed the data under the supervision of AT, EP, HAB and MM. HAB and MM jointly supervised the study. KN wrote the first manuscript draft, and MT, AT, EP, NMS, M-LHR, SQ, HAB and MM contributed with interpretation of results and reviewing and editing the manuscript.

## Data and code availability

Analysis code is available at: https://doi.org/10.5281/zenodo.21160475

The individual-level data underlying the results cannot be made publicly available due to data protection regulations. However, access to the data for researchers may be approved with some restrictions if necessary conditions are met.

## Conflict of interest

MM and SQ are cofounders and scientific advisors of Mirvie Inc. Mirvie had no role in study design, analysis, or result interpretation.

## Abbreviations

ACTA2: Smooth muscle α-actin
AFP: Alpha-fetoprotein
ANGPT2: Angiopoietin-2
CV: Cross-Validation
DAMP: Damage-Associated Molecular Pattern
EPS: Early Pregnancy Study
GA: Gestational Age
GAMM: Generalized Additive Mixed Model
GWAS: Genome Wide Association Study
ID: Identifier
IL-33: Interleukin 33
IL1RL1: Interleukin 1 receptor-like 1
ST2: sST2
IVW: Inverse Variance Weighted
LMOD1: Leiomodin-1
LMP: Last Menstrual Period
LOD: Limit of Detection
LOSO: Leave-One-Subject-Out
MAE: Mean Absolute Error
MR: Mendelian Randomization
NPX: Normalized Protein eXpression
PCA: Principal Component Analysis
pQTL: protein Quantitative Trait Loci
REML: REstricted Maximum Likelihood
RMSE: Root Mean Square Error
SD: Standard Deviation
SHAP: SHapley Additive exPlanations
UMAP: Uniform Manifold Approximation and Projection

## Notes

https://doi.org/10.5281/zenodo.21160475

