## Supplementary Note for "The molecular triggers of human labour: a longitudinal plasma proteomics study"

#### Table of Contents

**Supplementary Note 1: Comparing labour timing to published population data** **3**
Supplementary Table 1. Summary of delivery timing metrics. 3
Supplementary Figure 1. Kaplan–Meier curves for women not yet in labour from 38 weeks to delivery. 4
**Supplementary Note 2: Data exploration on quantitative proteomics data** **5**
Supplementary Figure 2. Scree plot from PCA analysis for the first 30 PCs. 5
Supplementary Figure 3. Projection of samples on the first two PCs. 6
Supplementary Figure 4. Parameter exploration for UMAP analysis. 7
Supplementary Figure 5. UMAP projection, as per Sup. Fig. 4. 7
Supplementary Figure 6. Parameter exploration for UMAP analysis. 8
Supplementary Figure 7. UMAP projection, as per Sup. Fig. 6. 8
**Supplementary Note 3: Longitudinal modeling with Generalized Additive Mixed Models (GAMMs)** **9**
Supplementary Figure 8. Results of the GAMMs analysis on the exploratory group. 10
Supplementary Figure 9. Results of the GAMMs analysis on the confirmatory group. 11
Supplementary Table 2. Proteins with significant pre-labour changes in both exploratory and confirmatory groups. 11
Supplementary Figure 10. Fitted smooth trajectories from each GAMM, produced by evaluating each model at 30 fixed time intervals, for both exploratory and confirmatory groups. 14
Supplementary Figure 11. Results of cross-validation for exploratory models. 16
**Supplementary Note 4: Gradient boosting model performance and benchmark comparisons in the exploratory group** **17**
Supplementary Figure 12. Bayesian optimization convergence plot. 17
Supplementary Table 3. Comparison of predictive models for days-to-delivery in the exploratory group. 17
Supplementary Figure 13. Bayesian optimization convergence plot for the XGBoost model with GA included. 18
**Supplementary Note 5: Gradient boosting model generalization and SHAP feature attribution in the confirmatory group** **19**
Supplementary Table 4. The top 20 features in the exploratory group, ranked by mean absolute SHAP values for the XGboost model that includes GA. 19
Supplementary Figure 14. Beeswarm plot for XGboost model with GA included on the exploratory and confirmatory group. 20
**Supplementary Note 6: Alternative interpretations and mechanistic considerations for the IL-33/ST2 axis findings** **22**
**Supplementary Note 7: Enrichment and Network analysis** **24**

Supplementary Figure 15. STRING ego network for AFP. 25
Supplementary Figure 16. STRING ego network for IL1RL1. 25
Supplementary Figure 17.  STRING ego network for ACTA2. 26
Supplementary Figure 18. STRING ego network for LMOD1. 27
Supplementary Figure 19. STRING ego network for ANGPT2. 27
**Supplementary Note 8: Comparison with prior work and replication dataset analysis on SomaScan 7k data 28**
Supplementary Figure 20. Core cascade proteins in GSE206454 replication dataset. 29
**Supplementary Note 9: Sensitivity analysis of cascade onset ordering 30**
Supplementary Figure 21. Cascade ordering sensitivity analysis: day of first detected change under three threshold definitions. 30
**Supplementary Note 10: Mendelian Randomization Analysis 32**
**References 35**

### Supplementary Note 1: Comparing labour timing to published population data

To place our population's timing of spontaneous labour in context, we compared its gestational-age distribution with the Early Pregnancy Study (EPS) reported by *Jukic et al.*[1]. EPS estimated gestational length from the day of ovulation (urinary hormones), reporting a median of 268 days and substantial natural variability even among healthy singleton pregnancies. Because our pipeline uses Last Menstrual Period (LMP) based gestational age (GA), we analyzed time-to-event from 38+0 weeks (266 days) to delivery using left-truncated Kaplan-Meier curves and treated spontaneous labour as the event (with acute C-sections censored). We summarized P(280) and P(287) as the probabilities of still being undelivered at 40 and 41 weeks, respectively, and reported the population median GA at delivery.

For interpretability against EPS's ovulation-based scale, we present two population reference curves. The LMP+14 alignment shifts the EPS ovulation-based distribution by 14 days – the conventional offset between ovulation and LMP – to place both datasets on a comparable scale. The spline-smoothed LMP variant used the smoothed LMP-based distribution directly reported by *Jukic et al.*, which provides a population reference for what a healthy singleton pregnancy distribution looks like on an LMP scale without assuming a fixed 14-day ovulation offset. Together these two references define the expected range of the population distribution and allow us to assess whether our population's delivery timing is representative of healthy term pregnancies despite its small size and selective inclusion criteria. Direct numerical comparisons are interpreted with the expected ~14-day offset between ovulation- and LMP-based timing.

In our exploratory group, the median LMP-based GA at delivery was 285 days for the labour definition. The probability of still being undelivered was P(280)=0.862 and P(287)=0.207. Under the LMP+14 alignment the median was 284 days with P(280)=0.635 and P(287)=0.323, while with spline-smoothed LMP the median was 285 days with P(280)=0.681 and P(287)=0.390. These survival estimates indicate that a substantial fraction of women remain pregnant through 40–41 weeks in our population, consistent with the broad late-gestation variability described by *Jukic et al.* (median 268 days on an ovulation-based scale, which is approximately 280 days on an LMP scale, and a naturally wide spread in healthy singleton pregnancies). Differences in medians and tail behaviour are expected given ovulation- vs LMP-based time origins, population composition and clinical practices, and our censoring of non-spontaneous deliveries.

**Supplementary Table 1. Summary of delivery timing metrics.**

For each analysis, the table reports the median GA at delivery (days) and the probabilities of still being undelivered at 40 weeks (P(280)) and 41 weeks (P(287)). Labour values come from our population, while LMP+14 and LMP spline are population benchmarks derived from *Jukic et al.*[1] on an LMP scale. All estimates are conditional on reaching 38 weeks. The event is spontaneous labour and acute caesarean deliveries are censored.

| Dataset | GA days (median) | P (280) | P (287) |
|---|---|---|---|
| labour | 285 | 0.862 | 0.207 |
| LMP+14 | 284 | 0.635 | 0.323 |
| LMP spline | 285 | 0.681 | 0.390 |

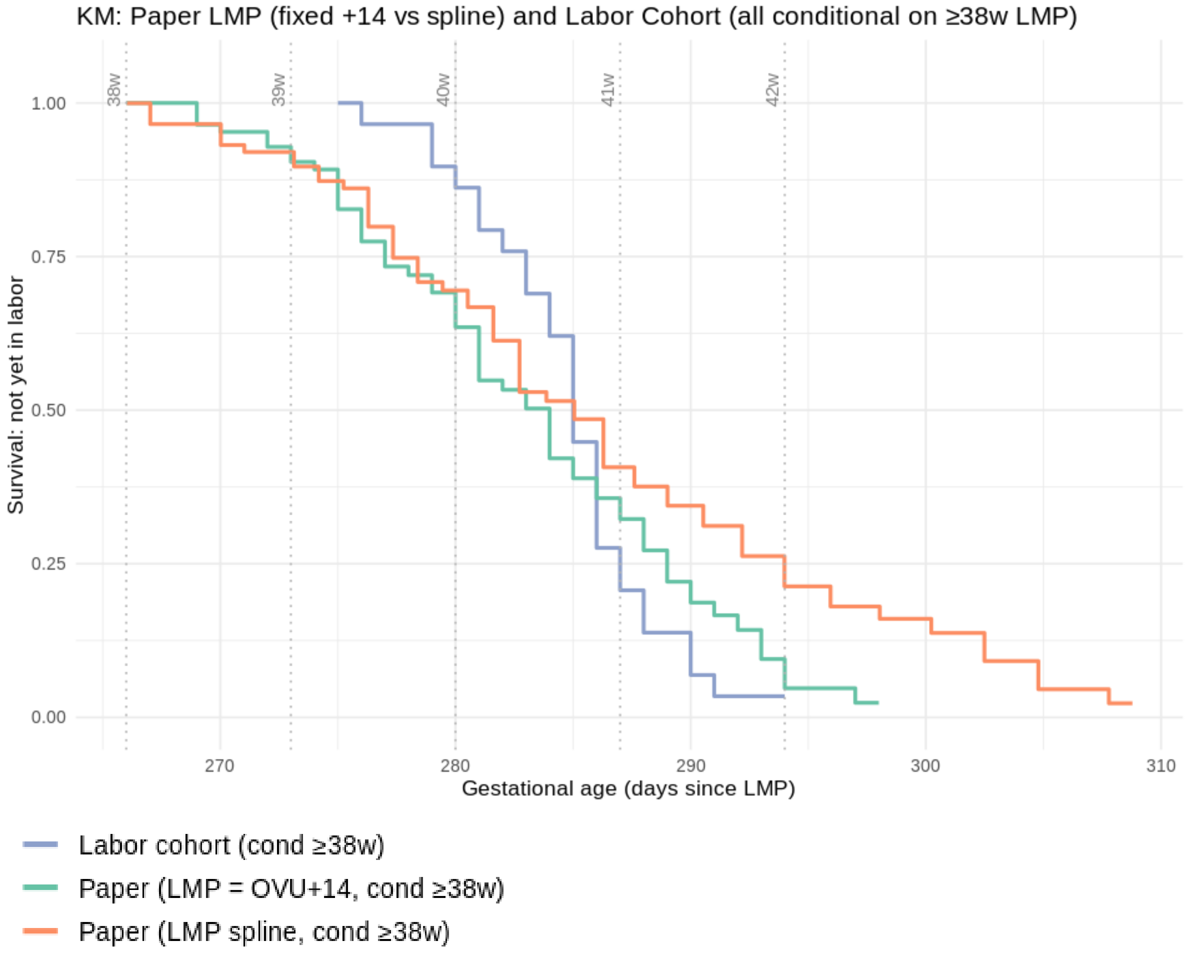


**Supplementary Figure 1. Kaplan–Meier curves for women not yet in labour from 38 weeks to delivery.**

The labour curve is our population (event = spontaneous labour; acute C-sections censored). The LMP+14 and LMP spline curves are population references reconstructed from Jukic et al.[1] mapped to an LMP scale: LMP + 14 shifts the ovulation-based distribution by 14 days. LMP spline uses the smoothed LMP-based distribution reported in that study. All curves are left-truncated at 266 days (38 weeks). Vertical dashed lines mark 280 days (40 w) and 287 days (41 w). The y-axis shows survival P(t) = probability not yet in labour, and the x-axis shows GA (days since LMP).

### Supplementary Note 2: Data exploration on quantitative proteomics data

The Olink Explore HT proteomics platform was used to perform quantitative proteomics analysis on our samples. Raw data was then processed and normalized, resulting in a final dataset of 5,416 proteins quantified for 491 samples split among 40 individuals at heterogeneous time points with respect to delivery date. As the full proteomics dataset was acquired as a single batch, we deemed batch effect correction unnecessary. We then performed preliminary data exploration to evaluate whether any structure was present in the data, using Principal Component Analysis (PCA) and Uniform Manifold Approximation and Projection (UMAP) to obtain a dimensionally-reduced representation of the data space.

We first performed a Principal Component Analysis (PCA) on the raw NPX values and considered the first two principal components (PCs). The scree plot indicated that the first two PCs explained 17% and 4.8% of the total variance, respectively (**Supplementary Figure 2**). We then visualized the projection onto these two components, colouring them according to time before delivery (**Supplementary Figure 3**). PCA projections showed a clear separation between pre-delivery and post-delivery samples, along PC2. Nonetheless, the first two PCs only accounted for 21.8% of the total variance.

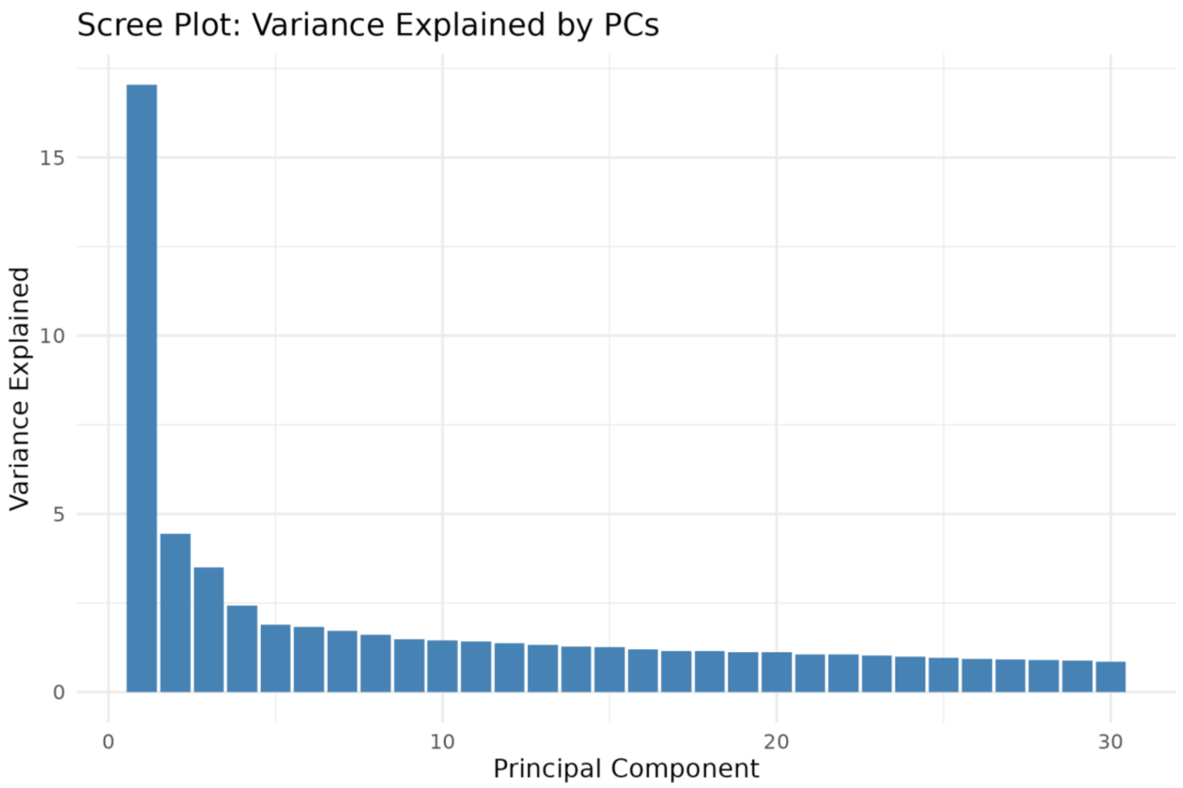


**Supplementary Figure 2. Scree plot from PCA analysis for the first 30 PCs.**

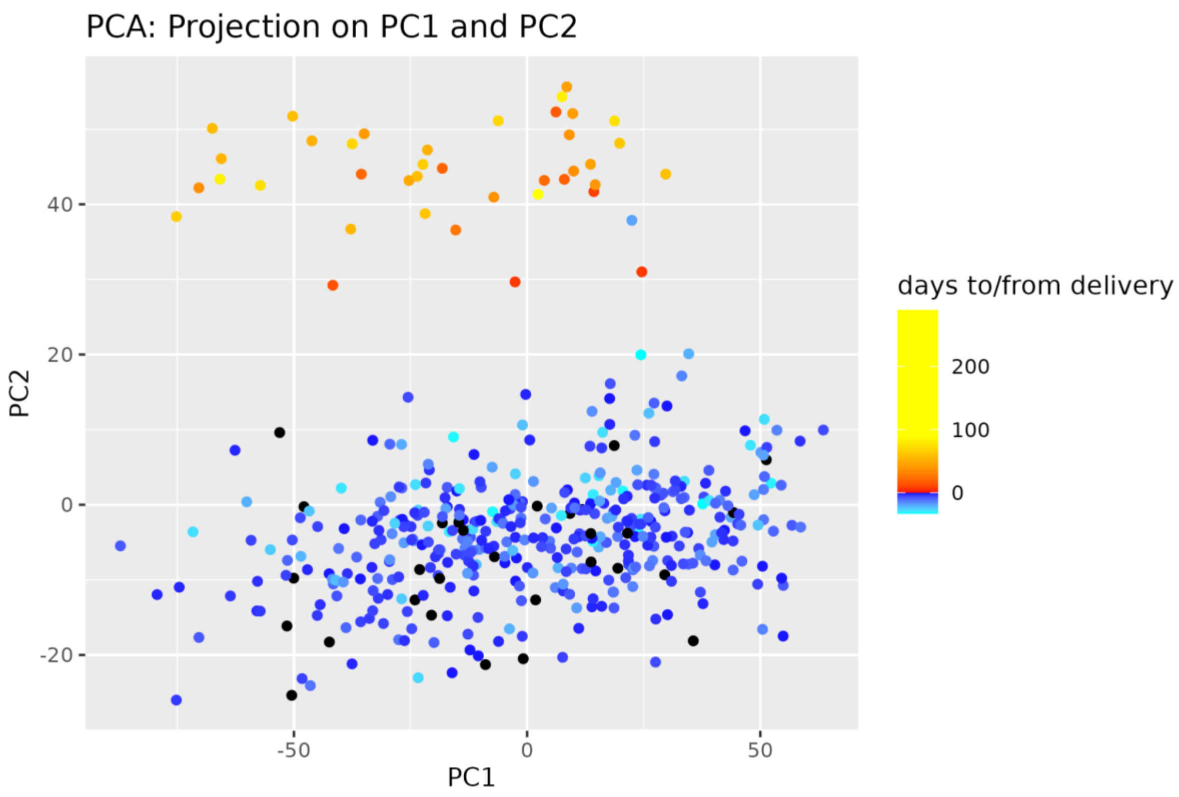


**Supplementary Figure 3. Projection of samples on the first two PCs.**

Samples are colored according to the associated number of days before delivery.

We then applied Uniform Manifold Approximation and Projection (UMAP) to the dataset, which provides a two-dimensional representation of the data while capturing both global structure and non-linear relationships between data points. To assess the robustness of the embedding, we performed hyperparameter exploration, varying the number of neighbours and minimum distance for the algorithm across discrete values. We therefore obtained multiple embeddings, which we coloured either by days before delivery (**Supplementary Figure 4 and 5**) or according to the identity of each individual participant they belonged to (**Supplementary Figure 6 and 7**). A representative UMAP projection using 20 neighbours and a minimum distance of 0.5 is shown in **Supplementary Figure 4** and **6**.

UMAP projections show no dependency on grouping with respect to time to delivery; rather, samples clustered on a per-individual participant basis, showing that differences between individuals determine the overall data structure more than pregnancy progression. Notably, our proteomics panel was not directly designed to study pregnancy or labour and includes a large number of proteins likely to be unrelated to the phenomenon under study, which drive data structure more than possibly subtle but important signals.

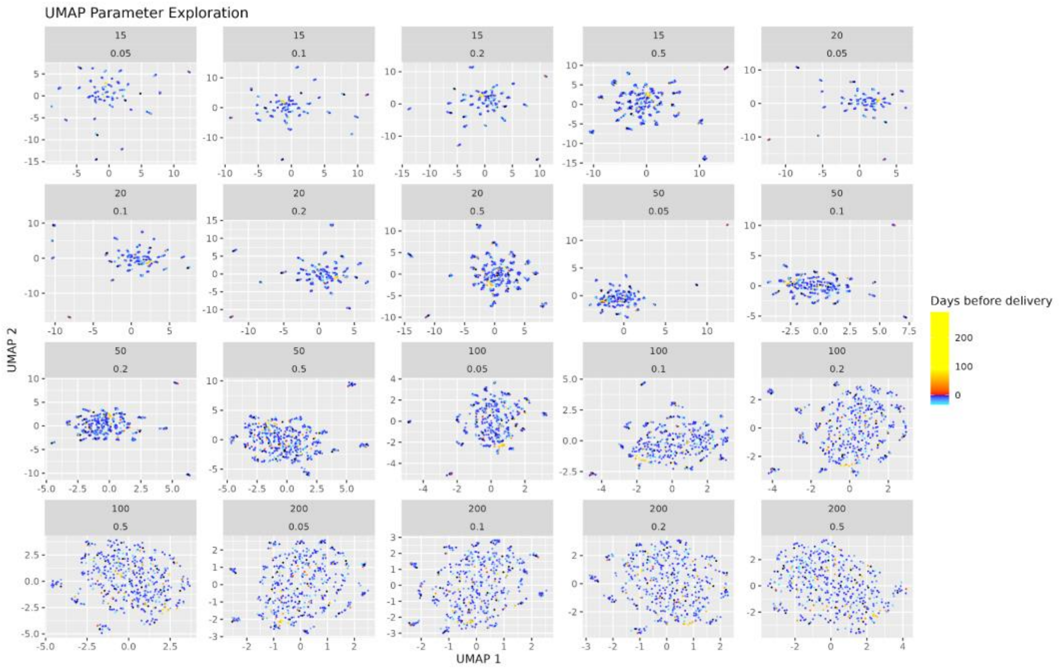


**Supplementary Figure 4. Parameter exploration for UMAP analysis.**

Each plot represents the UMAP projection of the proteomics data projected using a different value for number of neighbours (number at the top on the title of each plot) and minimum distance (number at the bottom). Each data point represents a sample of our dataset, and they have been coloured according to days before or after delivery.

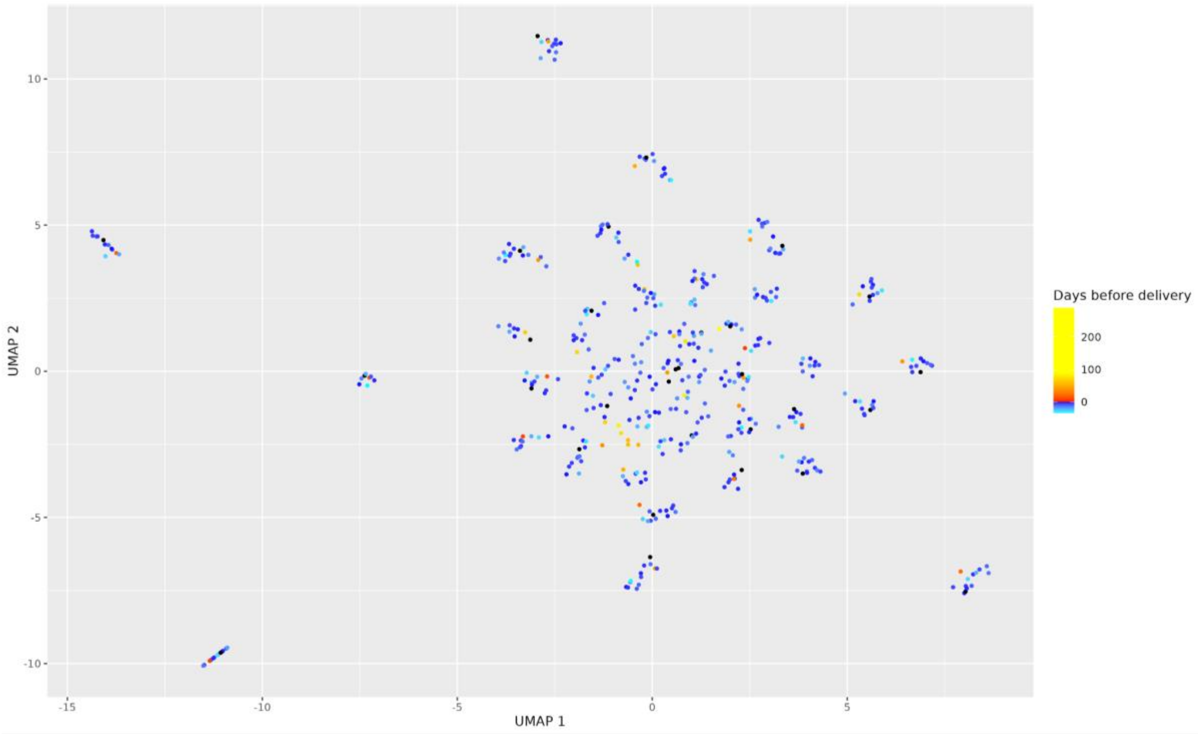


**Supplementary Figure 5. UMAP projection, as per Sup. Fig. 4.**

The projection is obtained using neighbours = 20 and minimum distance = 0.05.

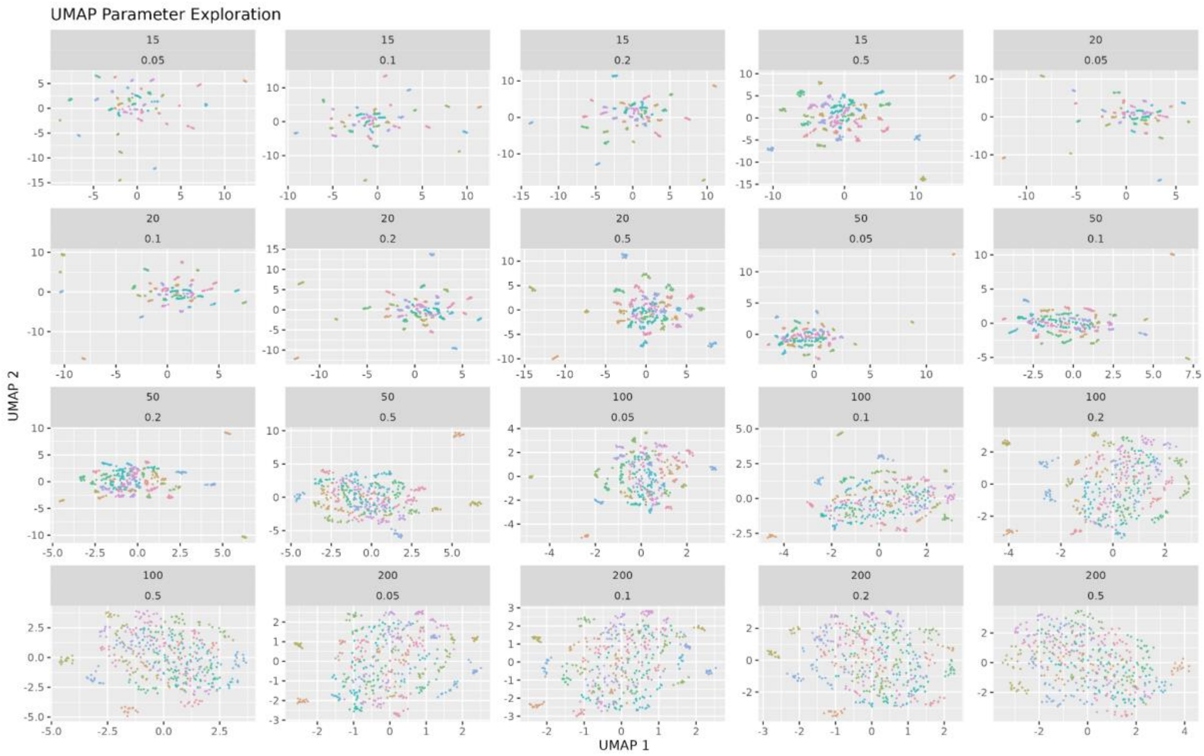


**Supplementary Figure 6. Parameter exploration for UMAP analysis.**

The analysis is done as per Sup. Fig. 4, except that each sample is coloured according to the individual it was taken from. As there are 40 individuals in the dataset, some of the colours end up being very similar.

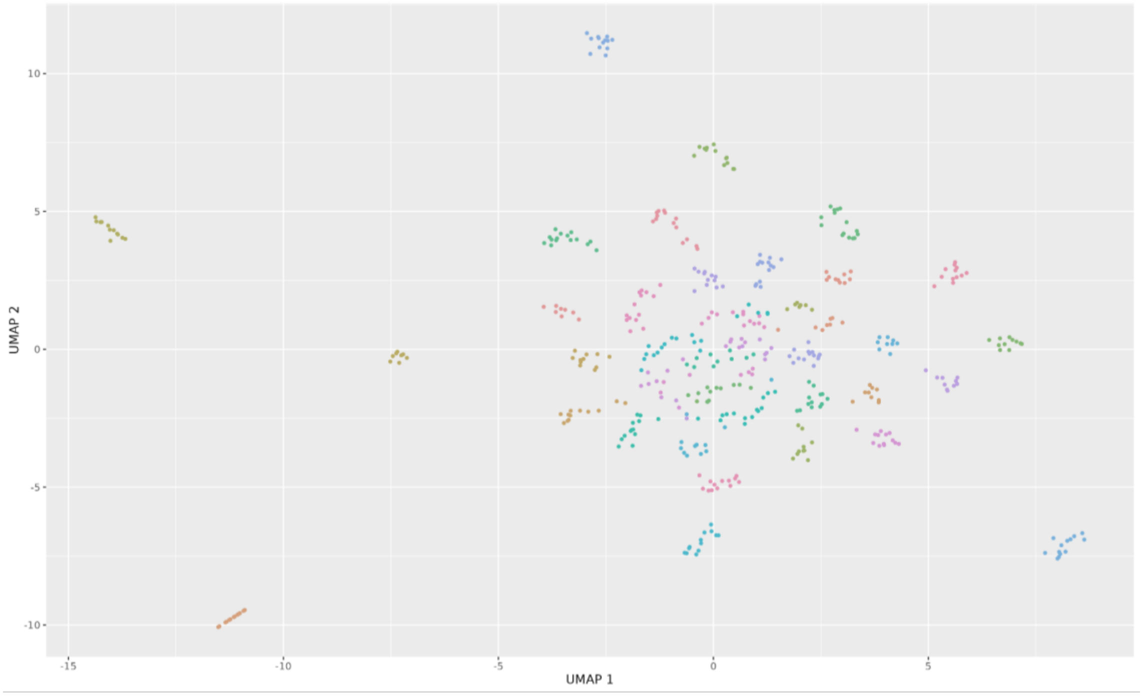


**Supplementary Figure 7. UMAP projection, as per Sup. Fig. 6.**

The projection is obtained using neighbours = 20 and minimum distance = 0.05.

### Supplementary Note 3: Longitudinal modeling with Generalized Additive Mixed Models (GAMMs)

We built a set of Generalized additive mixed models (GAMMs), fitting one model independently for each protein in the dataset. To make best use of the available data, samples were binned according to relevant time windows (see Methods) and only windows occurring before or at the delivery date were considered. We aimed at identifying proteins showing the most consistent temporal trends across individuals during the period of interest, and to characterize their behaviour (see Methods). We first performed the analysis on the exploratory group, using conditional $R^2$ ($cR^2$) as the primary measure of model quality, while also considering marginal $R^2$ ($mR^2$). $cR^2$ represents the proportion of variance explained by the full model, including both fixed and random effects, thereby capturing both temporal trends and variability between individuals, whereas $mR^2$ represents the proportion of variance explained by the fixed effects of the model alone, corresponding to a shared temporal trend between individuals.

**Supplementary Figure 8** showcases an overview of the models collected using the exploratory dataset. The distributions of conditional and marginal $R^2$ (**Supplementary Figures 8A and 8B**) differ markedly, with $mR^2$ strongly skewed towards lower values. This is consistent with our preliminary analysis, which indicated that differences between individuals are a major determinant of the overall data structure. These differences are captured by the random effects in the GAMMs and are therefore not reflected in $mR^2$. Nonetheless, several proteins achieved moderate $mR^2$ values, and these largely corresponded to the proteins with the highest $cR^2$ values in the dataset (**Supplementary Figure 8C**). On this basis, we focused our following efforts on a subset of 121 proteins having conditional $R^2 \geq 0.7$ (**Supplementary Figure 8D**).

We then applied the same approach to the confirmatory dataset (**Supplementary Figure 9**), overall resulting in a dataset with similar distributions of $cR^2$ and $mR^2$ as the exploratory dataset. However, a larger number of proteins reached $cR^2 \geq 0.7$ (276 over 121 of the exploratory dataset) and achieved larger maximum $mR^2$ values, also in this case corresponding to proteins with the largest $cR^2$ in the dataset.

The overlap between the exploratory and confirmatory datasets, only considering proteins with $cR^2 \geq 0.7$ in both cases, consisted of 28 proteins, including many of the top-scoring proteins found in the exploratory dataset (**Supplementary Table 2**).

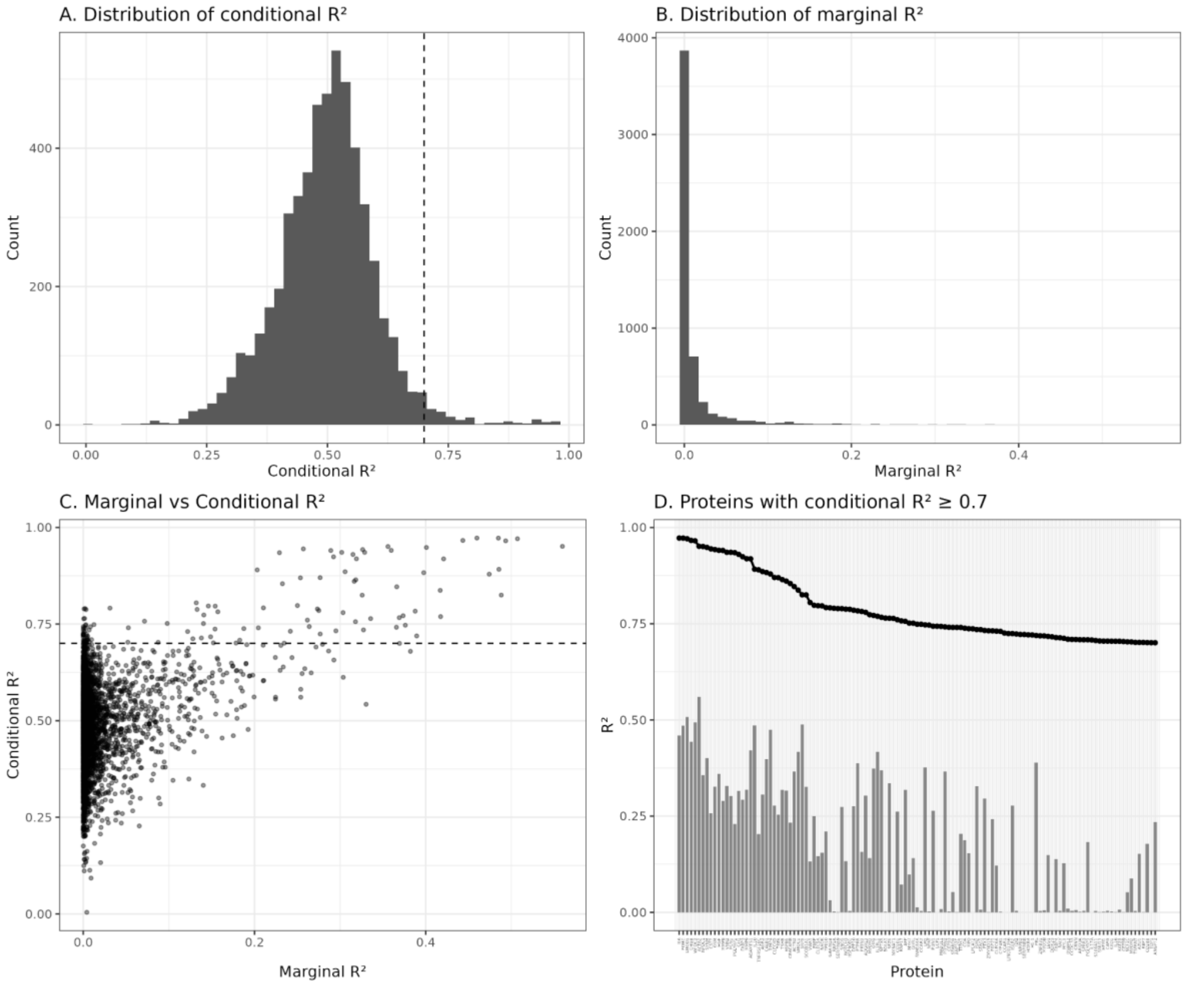


**Supplementary Figure 8. Results of the GAMMs analysis on the exploratory group.**

(A) distribution of conditional $R^2$ values over the whole dataset, (B) distribution of marginal $R^2$ values over the whole dataset, (C) Comparison of conditional and marginal $R^2$ values for each protein in the full dataset, (D) Conditional $R^2$ (black line) and marginal $R^2$ (grey bars) for each of the 121 proteins having Conditional $R^2 \geq 0.7$ in the dataset.

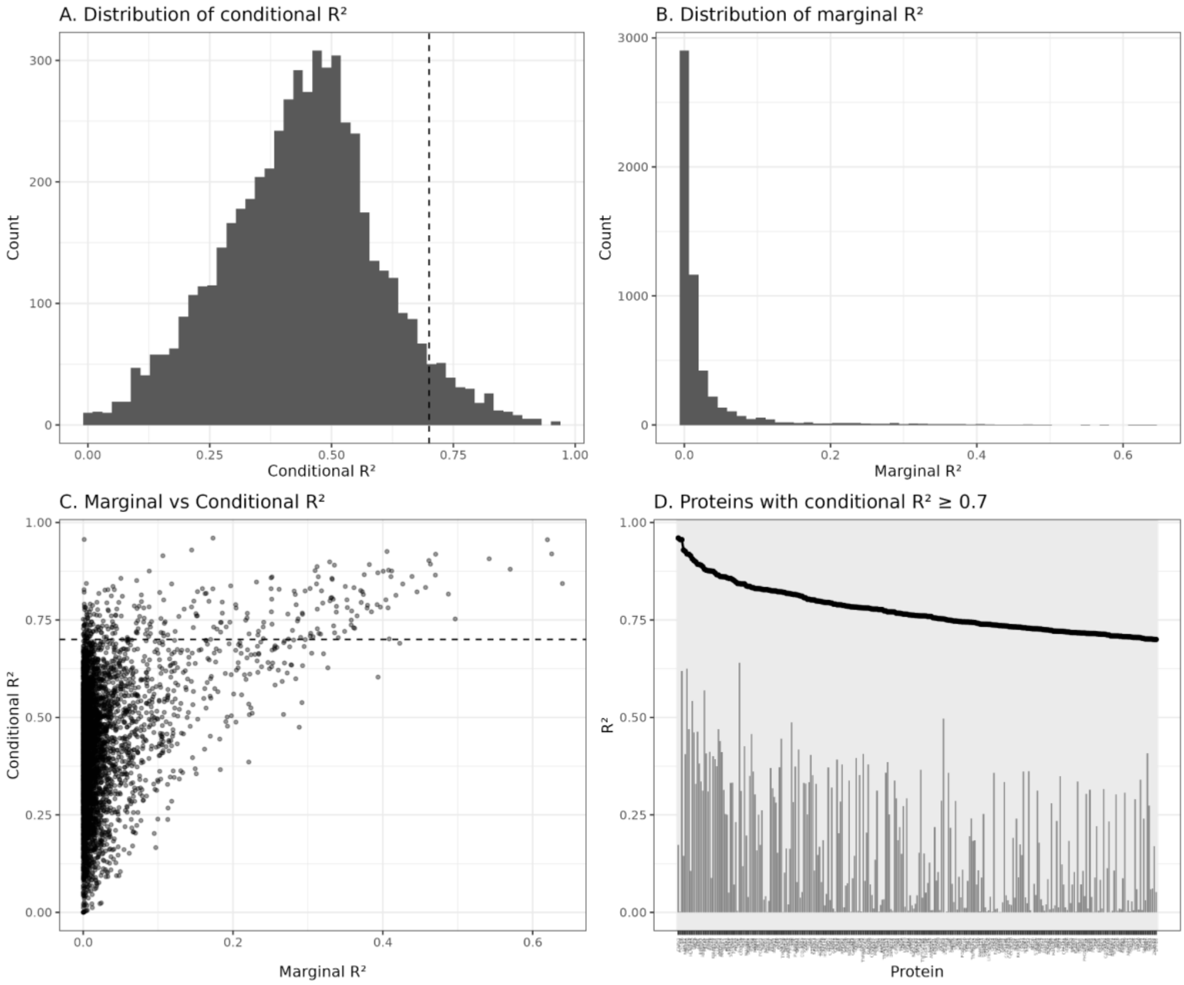


**Supplementary Figure 9. Results of the GAMMs analysis on the confirmatory group.**

(A) distribution of conditional $R^2$ values over the whole dataset, (B) distribution of marginal $R^2$ values over the whole dataset, (C) Comparison of conditional and marginal $R^2$ values for each protein in the full dataset, (D) Conditional $R^2$ (black line) and marginal $R^2$ (grey bars) for each of the 276 proteins having Conditional $R^2 \geq 0.7$ in the dataset.

**Supplementary Table 2. Proteins with significant pre-labour changes in both exploratory and confirmatory groups.**

Proteins with conditional $R^2 \geq 0.7$ in both the exploratory and confirmatory dataset. The rank columns refer to the position in the ranking, after sorting by conditional $R^2$ for decreasing values, for either dataset. Proteins are sorted by exploratory conditional $R^2$.

| Protein | UniProt AC | Exploratory $cR^2$ | Exploratory $mR^2$ | Exploratory rank | Confirmatory $cR^2$ | Confirmatory $mR^2$ | Confirmatory rank |
|---|---|---|---|---|---|---|---|
| IL6 | P05231 | 0.97 | 0.46 | 1 | 0.96 | 0.62 | 3 |
| POMC | P01189 | 0.97 | 0.48 | 2 | 0.7 | 0.41 | 270 |
| LMOD1 | P29536 | 0.97 | 0.51 | 3 | 0.92 | 0.63 | 6 |
| WFDC2 | Q14508 | 0.97 | 0.49 | 5 | 0.75 | 0.5 | 153 |

| Protein | UniProt AC | Exploratory $cR^2$ | Exploratory $mR^2$ | Exploratory rank | Confirmatory $cR^2$ | Confirmatory $mR^2$ | Confirmatory rank |
|---|---|---|---|---|---|---|---|
| ACTA2 | P62736 | 0.95 | 0.56 | 6 | 0.88 | 0.57 | 16 |
| NPDC1 | Q9NQX5 | 0.95 | 0.36 | 7 | 0.72 | 0.34 | 230 |
| SYNM | O15061 | 0.95 | 0.4 | 8 | 0.72 | 0.28 | 215 |
| OSM | P13725 | 0.94 | 0.33 | 10 | 0.78 | 0.35 | 105 |
| HGF | P14210 | 0.94 | 0.36 | 11 | 0.76 | 0.37 | 140 |
| MMP8 | P22894 | 0.94 | 0.29 | 12 | 0.88 | 0.41 | 19 |
| VNN2 | O95498 | 0.94 | 0.33 | 13 | 0.75 | 0.29 | 160 |
| FCN1 | O00602 | 0.94 | 0.3 | 14 | 0.89 | 0.38 | 13 |
| GH1 | P01241 | 0.93 | 0.31 | 16 | 0.8 | 0.4 | 77 |
| FABP3 | P05413 | 0.92 | 0.29 | 17 | 0.81 | 0.42 | 70 |
| MMP9 | P14780 | 0.92 | 0.32 | 18 | 0.71 | 0.31 | 257 |
| AGR2 | O95994 | 0.89 | 0.31 | 22 | 0.91 | 0.54 | 9 |
| CLEC4D | Q8WXI8 | 0.87 | 0.28 | 25 | 0.78 | 0.38 | 110 |
| TGFA | P01135 | 0.87 | 0.25 | 26 | 0.8 | 0.33 | 76 |
| HSPB6 | O14558 | 0.86 | 0.32 | 28 | 0.72 | 0.31 | 237 |
| PAEP | P09466 | 0.8 | 0.25 | 35 | 0.9 | 0.33 | 11 |
| DPP10 | Q8N608 | 0.79 | 0.28 | 45 | 0.71 | 0.01 | 265 |
| IL1RL1 | Q01638 | 0.77 | 0.42 | 51 | 0.84 | 0.64 | 36 |
| AFP | P02771 | 0.76 | 0.32 | 58 | 0.82 | 0.49 | 66 |
| ISM2 | Q6H9L7 | 0.73 | 0.33 | 76 | 0.71 | 0.22 | 241 |
| FGF21 | Q9NSA1 | 0.73 | 0.24 | 80 | 0.7 | 0.27 | 271 |
| FABP4 | P15090 | 0.71 | 0.14 | 96 | 0.73 | 0.31 | 207 |
| PGLYRP3 | Q96LB9 | 0.71 | 0.18 | 104 | 0.83 | 0.17 | 48 |
| ANGPT2 | O15123 | 0.7 | 0.23 | 121 | 0.87 | 0.47 | 24 |

We then evaluated whether the models generated from the exploratory and confirmatory datasets for these 28 overlapping high-quality proteins captured qualitatively similar time trends. To do so, we extracted fitted smooth trajectories from each GAMM by evaluating each model at 30 fixed time intervals and directly compared the predicted longitudinal patterns between the exploratory and confirmatory datasets (**Supplementary Figure 10**). This comparison shows that, in most cases, models derived from the two different datasets result in qualitatively similar trajectories (median Pearson's $r$ between exploratory and confirmatory datasets = 0.89). It is evident that some proteins display a largely linear and monotonic trend throughout the investigated period (ACTA2, AFP, ANGPT2, Il1RL1, ISM2, PGLYRP3), while others exhibit non-linear dynamics. Several proteins show a marked increase in abundance shortly before the onset of labour (AGR2, CLEC4D, FCN1, GH1, HGF, IL6, MMP8, MMP9, OSM, PAEP, TGFA, VNN2, WFDC2).

Among the 28 proteins with robust temporal trends identified by GAMM modelling, the five proteins with near-universal individual-level agreement across women (IL1RL1, ANGPT2, AFP, ACTA2, and LMOD1) are discussed in detail in the main manuscript. Here we focus on four proteins that show a sharp concordant rise concentrated in the final four days before delivery in both groups. The matrix metalloproteinases MMP8 and MMP9 show a sharp rise concentrated in the final four days before delivery in both groups (**Supplementary Figure 10**). The late rise is consistent with the well-established role of matrix metalloproteinases in cervical ripening[2,3], representing the activation of neutrophil- and macrophage-derived MMPs as part of the sterile inflammatory cascade[4,5] in the final days of parturition. Both interleukin 6 (IL-6) and Oncostatin M (OSM) show a sharp late rise in plasma levels concentrated in the final four days before delivery, with good cross-group concordance, a temporal pattern consistent with their shared biology as members of the IL-6 cytokine family signalling through the gp130 receptor complex[6]. IL-6 is a well-established mediator of the inflammatory transition to labour both in term[7] and preterm pregnancies[8]. IL-6 is stimulating prostaglandin synthesis, activating cervical stromal cells, and promoting myometrial contractility through JAK-STAT3 signalling[8,9], with elevated levels documented in amniotic fluid and cervicovaginal secretions in labour[10–13]. The late and acute nature of the IL-6 rise in our plasma proteomics data (concentrated in the final week rather than showing the gradual early-onset trajectory of IL1RL1) positions it as an acute inflammatory amplifier downstream of the upstream alarmin-initiated cascade rather than an initiating signal, consistent with the kinetics of acute-phase cytokine responses. OSM has established roles in decidual stromal cell maintenance[14]. Its co-rise with IL-6 in the final pre-delivery window suggests coordinated activation of the gp130 signalling axis as part of the terminal inflammatory priming programme. Moreover, OSM is known to upregulate MMP9 mRNA expression and activity in smooth muscle cells[15], providing a potential mechanistic link between the late gp130 cytokine activation and the terminal MMP-driven cervical remodelling programme, and suggesting that the convergent late rise of OSM, IL-6, MMP8 and MMP9 reflects a coordinated inflammatory effector programme rather than four independent signals.

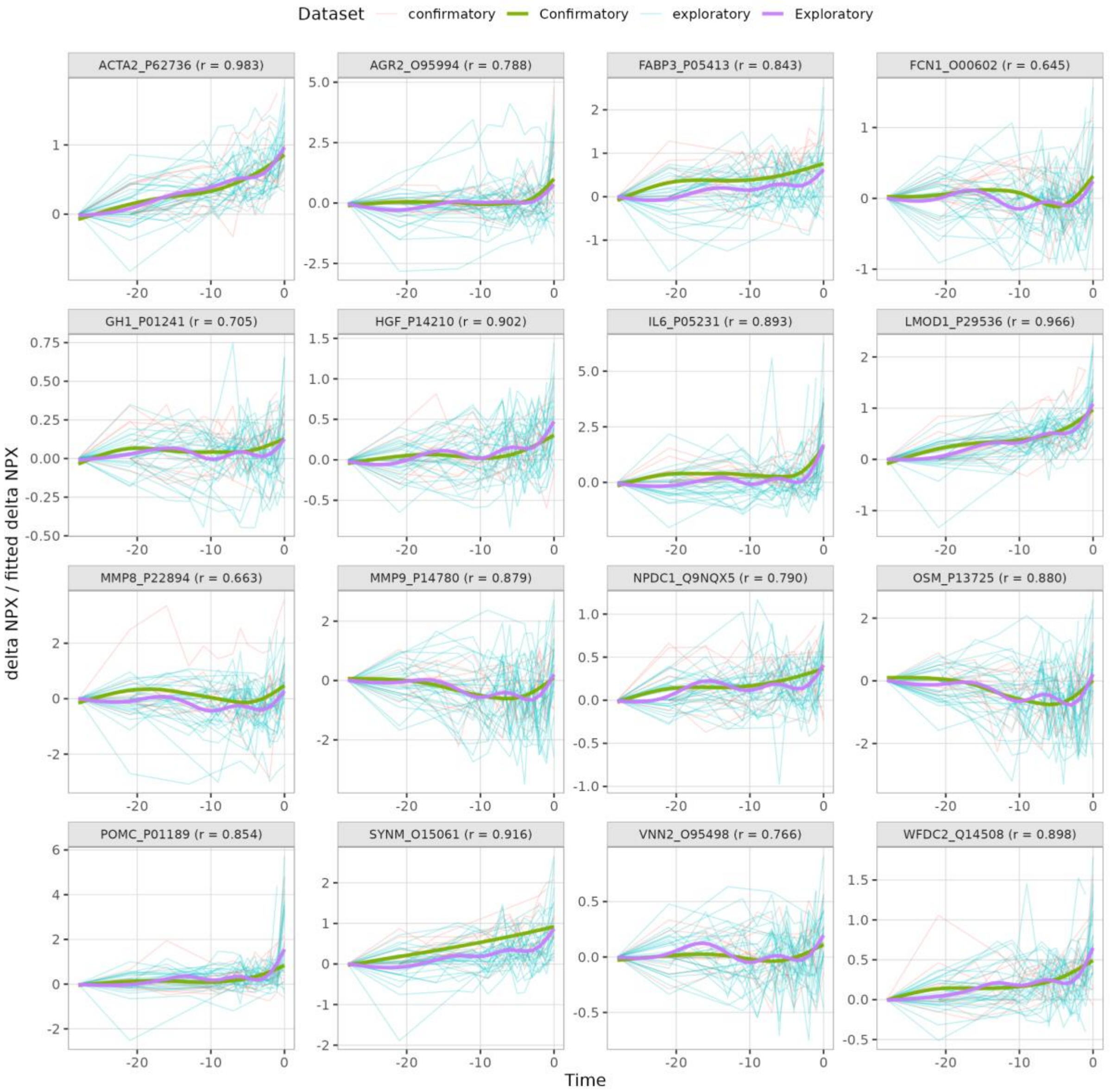


**Supplementary Figure 10. Fitted smooth trajectories from each GAMM, produced by evaluating each model at 30 fixed time intervals, for both exploratory and confirmatory groups.**

These plots are for 16 out of the 28 proteins and include both model fit and individual data trajectories. Pearson correlation coefficient between the confirmatory and exploratory models is indicated in parenthesis. Thin lines represent individual womens' observed ΔNPX trajectories and bold lines represent population-level GAMM smooths. Teal = exploratory group (n=30), purple = exploratory GAMM smooth, orange = confirmatory group (n=10), green = confirmatory GAMM smooth.

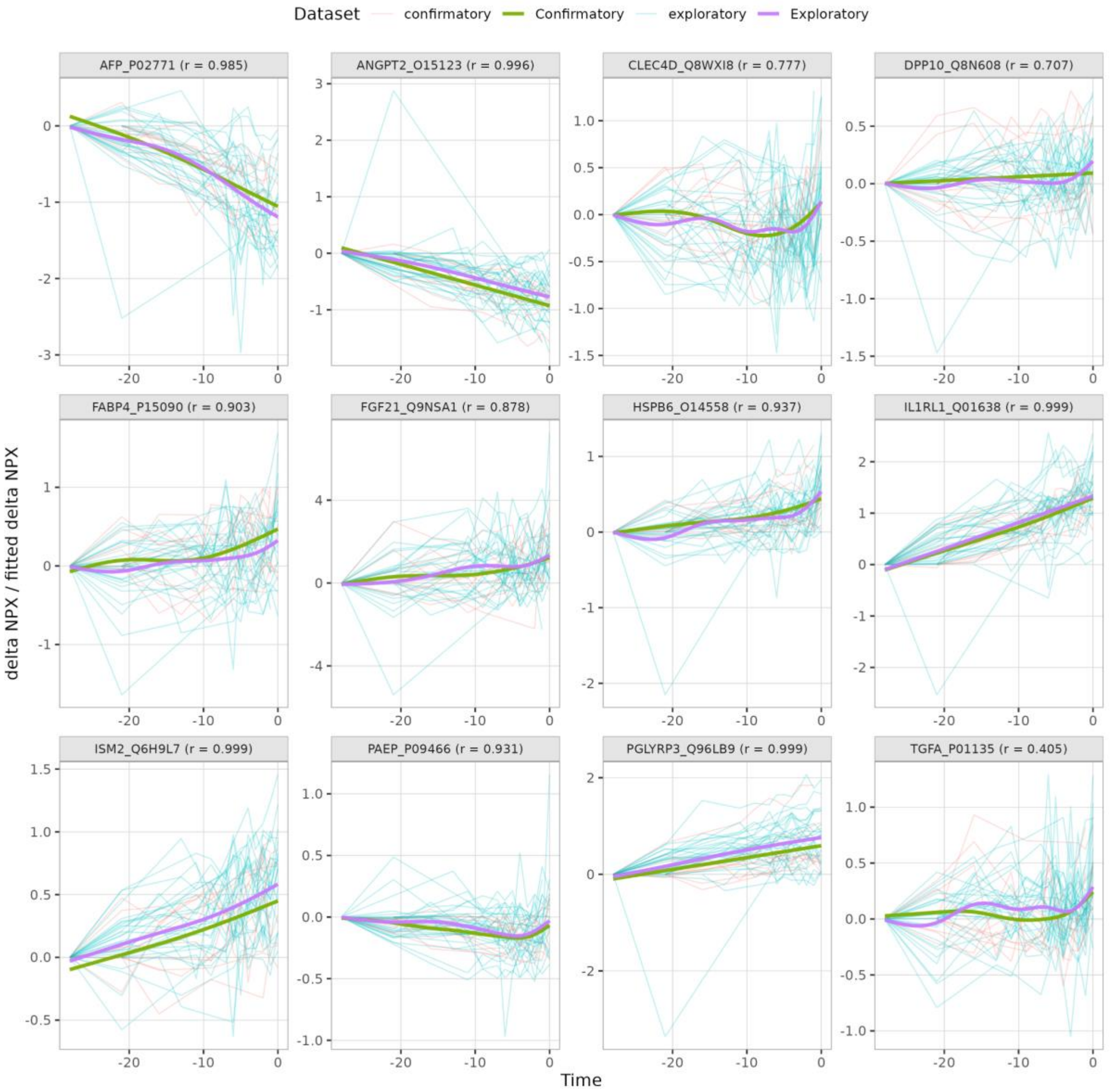


**Supplementary Figure 10 (continued). Fitted smooth trajectories from each GAMM, produced by evaluating each model at 30 fixed time intervals, for both exploratory and confirmatory groups.**
These plots are for the remaining 12 out of the 28 proteins. Pearson correlation coefficient between the confirmatory and exploratory models is indicated in parenthesis. Thin lines represent individual womens' observed ΔNPX trajectories and bold lines represent population-level GAMM smooths. Teal = exploratory group (n=30), purple = exploratory GAMM smooth, orange = confirmatory group (n=10), green = confirmatory GAMM smooth.

Finally, to assess the robustness and generalizability of our models, we performed person-level 5-fold cross-validation on the exploratory group. Individuals were partitioned into 5 groups, and for each fold, models were trained on a subgroup of individuals and evaluated on held-out individuals. Prediction error was quantified using the root mean squared error (RMSE). We then examined the distribution of both the observed NPX differences and the cross-validated RMSE values across proteins (**Supplementary Figure 11**). Prediction errors were found to be generally low and narrowly distributed, with RMSE values smaller than the observed variability in NPX measurements. This indicates that the protein models generalize well to unseen individuals. Variability across proteins also showcases the heterogeneous quality of our models.

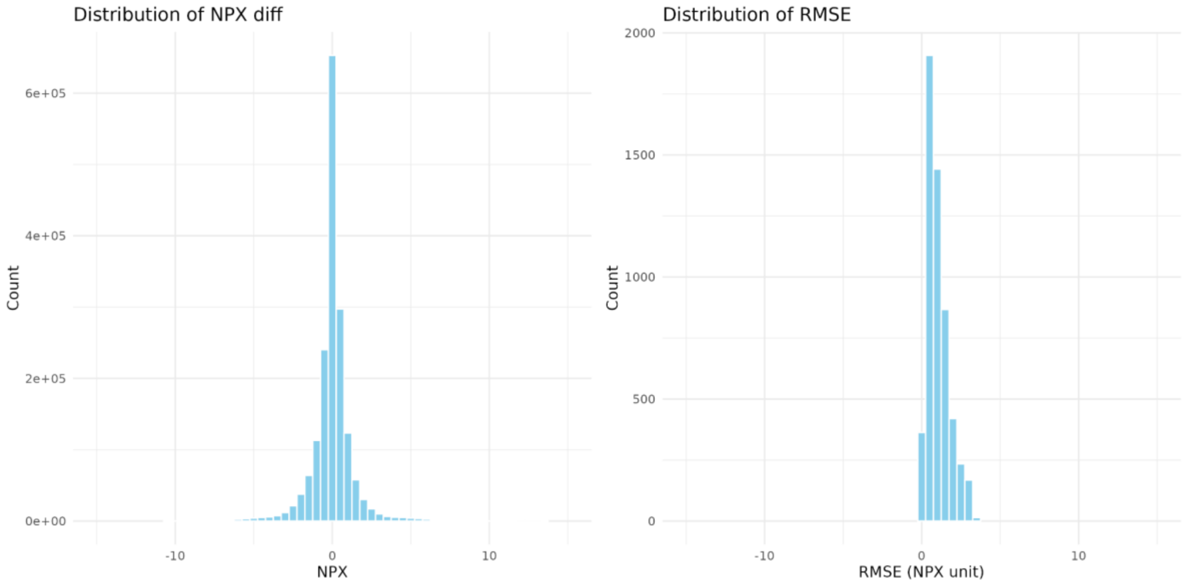


**Supplementary Figure 11. Results of cross-validation for exploratory models.**

Left: Distribution of NPX difference data in the dataset; right: distribution of RMSE after cross-validation for our dataset.

### Supplementary Note 4: Gradient boosting model performance and benchmark comparisons in the exploratory group

Bayesian optimization over XGBoost hyperparameters converged to a stable set of parameters at iteration 48 (learning rate = 0.01, maximum depth = 3, subsample = 0.5, colsample_bytree = 0.5, min_child_weight = 10.0, λ = 0.09, α = 0.01). The optimization procedure was performed with 10 random initialization points and 40 acquisition rounds (**Supplementary Figure 12**).

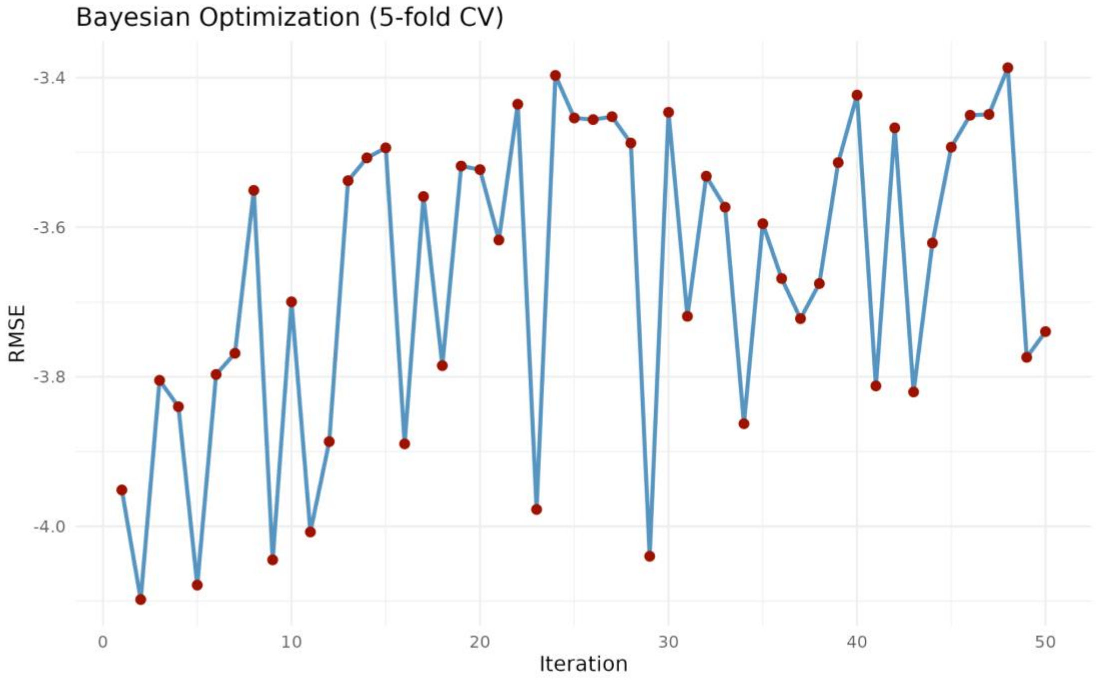


**Supplementary Figure 12. Bayesian optimization convergence plot.**

XGBoost gradient boosting trained on the full proteomic feature set achieved a mean Leave-One-Subject-Out (LOSO) Root Mean Square Error (RMSE) of 3.05 ± 1.05 days and Mean Absolute Error (MAE) of 2.45 ± 1.04 days in the exploratory group, outperforming all benchmarks including Random Forest (RMSE 4.79 ± 0.75), Lasso (RMSE 3.60 ± 2.06), Elastic Net (RMSE 3.89 ± 2.44), and a deterministic baseline assuming delivery at 40+0 weeks (RMSE 6.43 days respectively; **Supplementary Table 3**). LOSO CV was used rather than standard k-fold to prevent data leakage, since multiple samples from the same woman are temporally correlated, and including samples from the same woman in both training and validation folds would artificially inflate performance estimates.

**Supplementary Table 3. Comparison of predictive models for days-to-delivery in the exploratory group.**

LOSO Cross-validation (CV) performance of machine learning models and deterministic baselines predicting absolute days from delivery at the time of sampling, in the exploratory group (n=30 women). RMSE and MAE are reported in days. For learned models (XGBoost, Lasso, Elastic Net, Random Forest), values are mean ± standard deviation across LOSO folds. For a deterministic baseline (assuming delivery at exactly 40+0 weeks of gestation) a single RMSE is reported computed across all available samples; MAE is not reported for the baseline as predicted values are fixed. XGBoost was the best-performing proteomics-only model, outperforming all alternative learned models and both deterministic baselines. Lasso and Elastic Net showed high fold-to-fold variability, likely reflecting instability of linear regularization under the high-dimensional sparse feature space.

| Model | Mean RMSE ± sd | Mean MAE ± sd |
|---|---|---|
| XGBoost | 3.05 ± 1.05 | 2.45 ± 1.04 |
| Lasso | 3.60 ± 2.06 | 3.20 ± 2.01 |
| Elastic Net | 3.89 ± 2.44 | 3.41 ± 2.28 |
| Random Forest | 4.79 ± 0.75 | 3.89 ± 0.78 |
| Baseline 40+0 | 6.43 | |

With GA included, Bayesian optimization identified a configuration at iteration 33 (learning rate = 0.01, maximum depth = 3, subsample = 0.6, colsample_bytree = 0.7, min_child_weight = 7, $\lambda = 1.7$, $\alpha = 0.01$) with a grouped 5-fold CV mean RMSE of 2.85 days (**Supplementary Figure 13**). Using these settings in LOSO CV on the exploratory group yielded a mean RMSE = 2.60 ± 1.75 days and MAE = 2.18 ± 1.60 days.

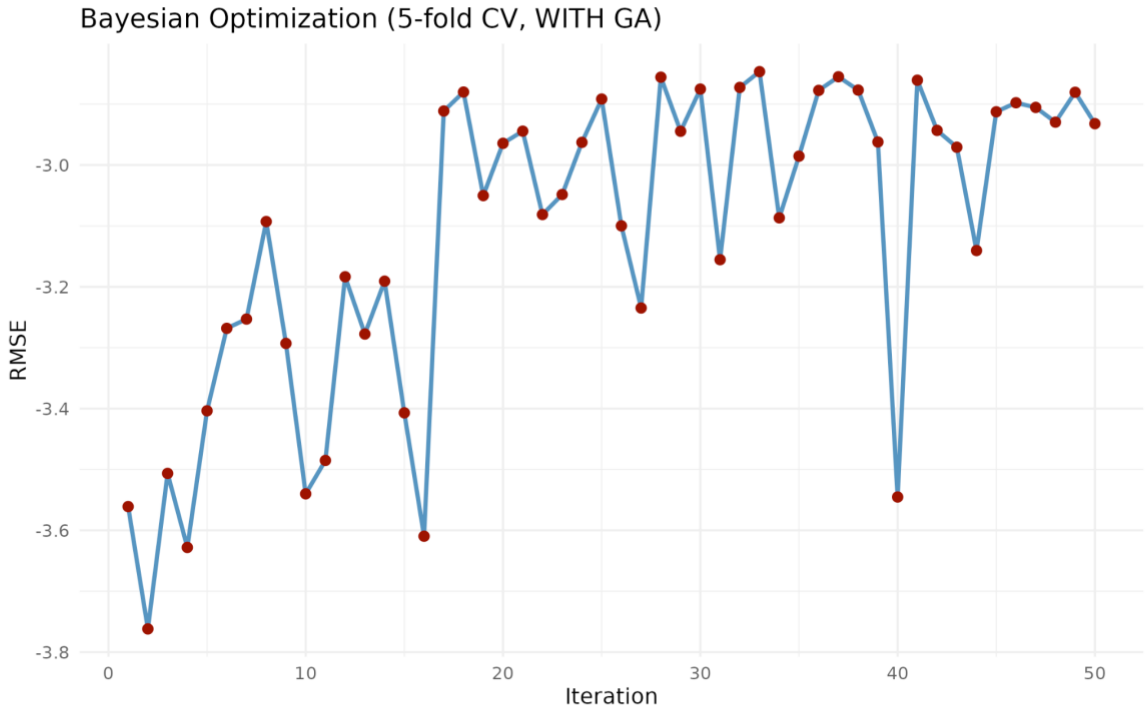


**Supplementary Figure 13. Bayesian optimization convergence plot for the XGBoost model with GA included.**

### Supplementary Note 5: Gradient boosting model generalization and SHAP feature attribution in the confirmatory group

SHAP (SHapley Additive exPlanations)[16] is a framework for interpreting machine learning model predictions by decomposing each prediction into additive contributions from individual features. For each sample, the SHAP value of a given protein quantifies how much that protein's level at that timepoint pushed the model's prediction of days-to-delivery above or below the baseline prediction. A positive SHAP value means the feature contributed toward predicting more days remaining until delivery; a negative SHAP value means it contributed toward predicting imminent delivery. Global feature importance is summarised as the mean absolute SHAP value across all samples, reflecting each protein's average contribution to predictions regardless of direction. Because SHAP values are sample-level, they also capture temporal structure: a protein that changes gradually over four weeks will have its predictive contribution distributed across many timepoints, whereas a protein that changes sharply only near delivery will show concentrated high-SHAP values in the final days.

LOSO CV on the confirmatory group yielded a mean RMSE of 3.70 days and MAE of 2.8 days, with 16 of the top 20 SHAP-ranked proteins overlapping between groups, including all top 5 (**Supplementary Table 4, Supplementary Figure 14**). Gestational age at sampling was the strongest single predictor in both groups, reflecting the well-established relationship between proximity to term and labour onset. The protein features identified here therefore capture biological variation in labour timing beyond what gestational age alone explains, which is the clinically and biologically relevant signal for understanding individual differences in the timing of spontaneous labour.

**Supplementary Table 4. The top 20 features in the exploratory group, ranked by mean absolute SHAP values for the XGboost model that includes GA.**

| Feature | UniProt AC | SHAP value exploratory | SHAP value confirmatory |
|---|---|---|---|
| Gestational age | – | 3.30 | 3.45 |
| AFP | P02771 | 0.74 | 0.63 |
| ANGPT2 | O15123 | 0.70 | 0.70 |
| ACTA2 | P62736 | 0.60 | 0.66 |
| LMOD1 | P29536 | 0.59 | 0.59 |
| IL1RL1 | Q01638 | 0.23 | 0.23 |
| KRT19 | P08727 | 0.19 | 0.21 |
| STC1 | P52823 | 0.15 | 0.15 |
| POLR3F | Q9H1D9 | 0.05 | 0.04 |
| ALPP | P05187 | 0.05 | 0.05 |

| | | | |
|---|---|---|---|
| UBR7 | Q8N806 | 0.05 | 0.03 |
| RNF4 | P78317 | 0.04 | 0.05 |
| C10orf95 | Q9H7T3 | 0.04 | 0.04 |
| CUX2 | O14529 | 0.04 | 0.04 |
| TIE1 | P35590 | 0.04 | 0.03 |
| SLITRK6 | Q9H5Y7 | 0.04 | 0.04 |
| DHPS | P49366 | 0.03 | 0.04 |
| FSTL3 | O95633 | 0.03 | 0.04 |
| FAM131B | Q86XD5 | 0.03 | 0.04 |
| FCN2 | Q15485 | 0.03 | 0.04 |

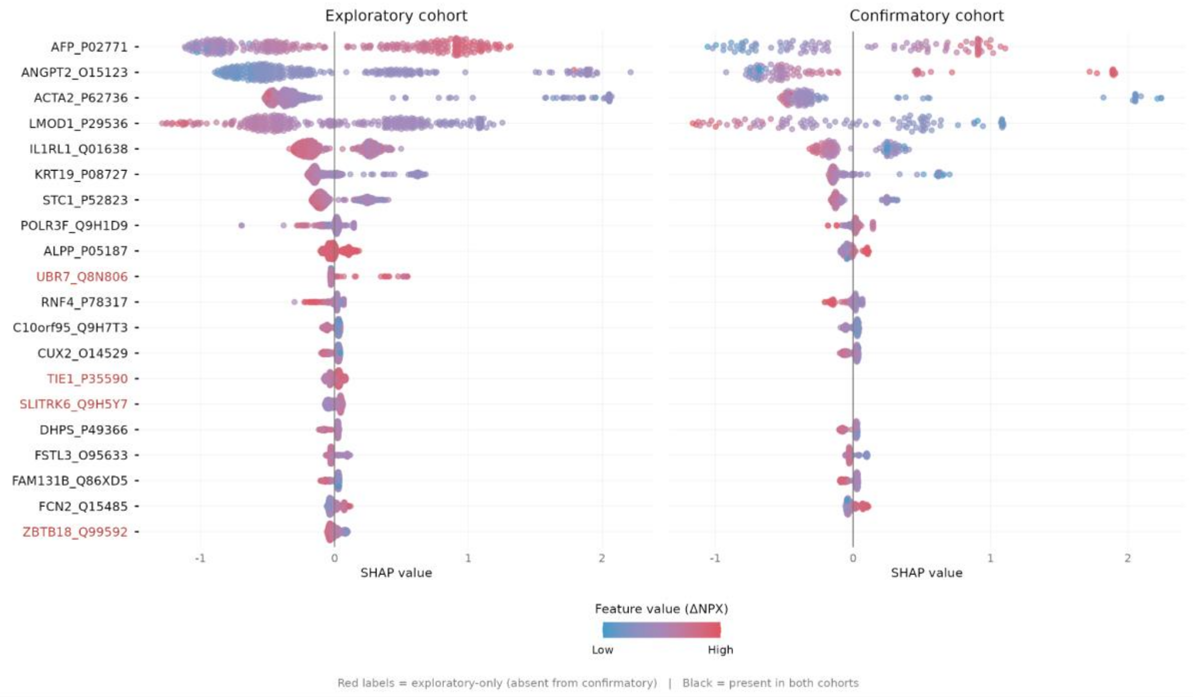


**Supplementary Figure 14. Beeswarm plot for XGboost model with GA included on the exploratory and confirmatory group.**

The top 20 proteins based on mean absolute SHAP values in the exploratory group are shown. 16 out of the top 20 proteins are overlapping in the exploratory and confirmatory groups. AFP shows the strongest and most consistent signal across both groups, with declining levels (blue points) driving positive SHAP contributions, meaning falling AFP is the model's strongest single push toward predicting imminent delivery. ANGPT2 follows a similar directional logic, with its dense cloud of blue points on the negative SHAP side. ACTA2 and LMOD1 show the mirror pattern, i.e. rising levels (red points) associate with positive SHAP contributions. IL1RL1's SHAP distribution appears modest in this global summary, with points clustering tightly near zero across both groups. GAMM modelling (**Supplementary Figure 10**) shows that IL1RL1 rises early and monotonically over the full four-week window, so its predictive contribution is distributed across many time points rather than concentrated at any single one, and the global mean |SHAP| systematically underrepresents features whose importance is temporally diffuse rather than window-specific.

### Supplementary Note 6: Alternative interpretations and mechanistic considerations for the IL-33/ST2 axis findings

IL-33 (O95760) was undetectable in peripheral plasma in 490 of 491 pre-partum samples across both groups (99.8%), compared to IL1RL1 (Q01638) which was detected in 100% of samples. Limits of detection (LOD) were calculated from negative control samples included on each plate using the negative control LOD method (NCLOD) as implemented in the OlinkAnalyze R package[17], whereby the LOD for each assay is derived from the distribution of negative control measurements within each plate. A sample was classified as above the LOD if its NPX value exceeded the plate-specific NCLOD threshold for that assay.

Bessa et al. (2014) demonstrated how the nuclear localisation of IL-33 is itself an active containment mechanism. Abolishing the nuclear localisation signal causes constitutive IL-33 release into serum and lethal systemic inflammation, establishing that nuclear retention normally prevents IL-33 from reaching the circulation even when it is being produced[18]. The absence of detectable plasma IL-33 in our population is therefore consistent with physiological nuclear containment being maintained throughout the pre-labour period, with only small quantities of IL-33 escaping into the extracellular space during tissue stress, producing quantities sufficient for local paracrine ST2L signaling[19] but insufficient for peripheral plasma accumulation, consistent with IL-33 acting as a local co-inducer of rising sST2 as part of escalating pathway activation rather than accumulating systemically[20]. The concentrations required for autocrine or paracrine ST2L activation are expected to be substantially lower than those required for peripheral plasma accumulation, making the plasma compartment an inherently insensitive readout of local IL-33 activity.

An important alternative interpretation of rising sST2 is that it reflects a hepatic acute-phase response independent of IL-33 signaling, as has been documented in preeclampsia and other inflammatory states where sST2 is produced by the liver in response to IL-6 and other acute-phase cytokines[21,22]. Several features of our data argue against this as the primary explanation in the present context. First, our population comprises healthy women with uncomplicated term pregnancies without evidence of infection or preeclampsia, in whom a sustained hepatic acute-phase response would be biologically unexpected. Second, IL-6 – the principal driver of hepatic acute-phase sST2 production – appears to be rising only towards the last days before labour in our GAMM analysis (**Supplementary Figure 10**), whereas IL1RL1 rises from the earliest sampling timepoints, making it implausible that the early IL1RL1 rise is driven by IL-6-mediated hepatic signaling. Third, the gradual, monotonic trajectory of IL1RL1 over four weeks is more consistent with a sustained biological program, such as progressive IL-33 release from senescent reproductive tissues, than with the typically transient and reactive kinetics of an acute-phase response. Together, these observations support progressive IL-33/ST2 pathway activation – rather than a hepatic acute-phase response – as the more parsimonious explanation for the early and sustained rise in sST2 in the specific context of spontaneous term labour.

We acknowledge that rising plasma sST2 cannot in isolation establish which direction the sST2:ST2L balance is shifting at the tissue level – the functional outcome depends on the relative expression of both isoforms in reproductive tissues, which peripheral plasma measurement cannot resolve. However, the consistent absence of detectable plasma IL-33 even in the last days before delivery is compatible with both a sequestration interpretation (sST2 capturing locally released IL-33 before it can signal) and an activation interpretation (IL-33 co-inducing sST2 as pathway activity escalates, with local concentrations remaining below the threshold for systemic detection regardless of signalling state). The Mendelian randomisation result (genetically higher IL1RL1 levels are associated with shorter rather than longer gestational duration) is more consistent with the activation interpretation, under which a higher IL-33/ST2 signalling tone accelerates the parturition programme, than with the sequestration interpretation, under which higher sST2 should prolong quiescence and delay delivery. Regardless of which interpretation is ultimately correct, the consistent and temporally specific rise of plasma sST2 as the earliest and most reproducible signal in our population, and its independent predictive contribution across a >5,000-protein panel, constitute empirical evidence that the IL-33/ST2 axis is engaged in the biological program preceding spontaneous labour onset.

### Supplementary Note 7: Enrichment and Network analysis

We analyzed the interaction context of the five overlapping proteins from GAMM and XGboost modeling in Cytoscape (v3.10.4)[23] and stringApp (v2.2.0)[24] querying the STRING database of molecular interactions (v12.0)[25]. We constructed ego networks for each of these proteins to characterize their local interaction neighborhoods. STRING queries were restricted to *Homo sapiens* (taxonID: 9606) with a combined score cutoff of 0.90 (highest confidence) and up to 20 direct interaction partners. All network construction, enrichment analysis, and visualisation were performed using R (v4.3) with the RCy3 package[26] (v2.24.0) interfacing with Cytoscape via the CyREST API.

For each protein set, we ran functional enrichment analysis in STRING against a whole-genome background, using the stringApp built-in enrichment functionality. We queried Gene Ontology annotations[27] (Molecular Function, Biological Process, Cellular Component), UniProt keywords[28], KEGG pathways[29], Reactome pathways[30], WikiPathways[31], Monarch human phenotypes[32], Pfam[33] and SMART[34] protein domains, InterPro protein features[35], local STRING clusters[36], DISEASES[37], COMPARTMENTS[38], and TISSUES[39], which integrates tissue expression evidence from the Human Protein Atlas (HPA)[40], GTEx[41], and other sources. Multiple testing was controlled with Benjamini–Hochberg False Discovery Rate (FDR), reporting terms with FDR≤0.05, which were considered significant. Full results of the analysis are presented in **Supplementary Table 5**, which is provided as a separate Excel file (*Supplementary_Table_5.xlsx*) due to its size.

Redundant enrichment terms were filtered using a Jaccard similarity cutoff of 0.9, and up to five non-redundant terms per network were selected for visualisation, excluding UniProt Keywords and STRING Cluster annotations. Selected enrichment terms were displayed as pie chart overlays on network nodes using the stringApp chart functionality, with each slice representing one enrichment term. A colour-coded legend indicating the selected terms and their associated FDR values was added as a network annotation.

The ego networks for the 5 proteins are shown in **Supplementary Figures 15-19**.

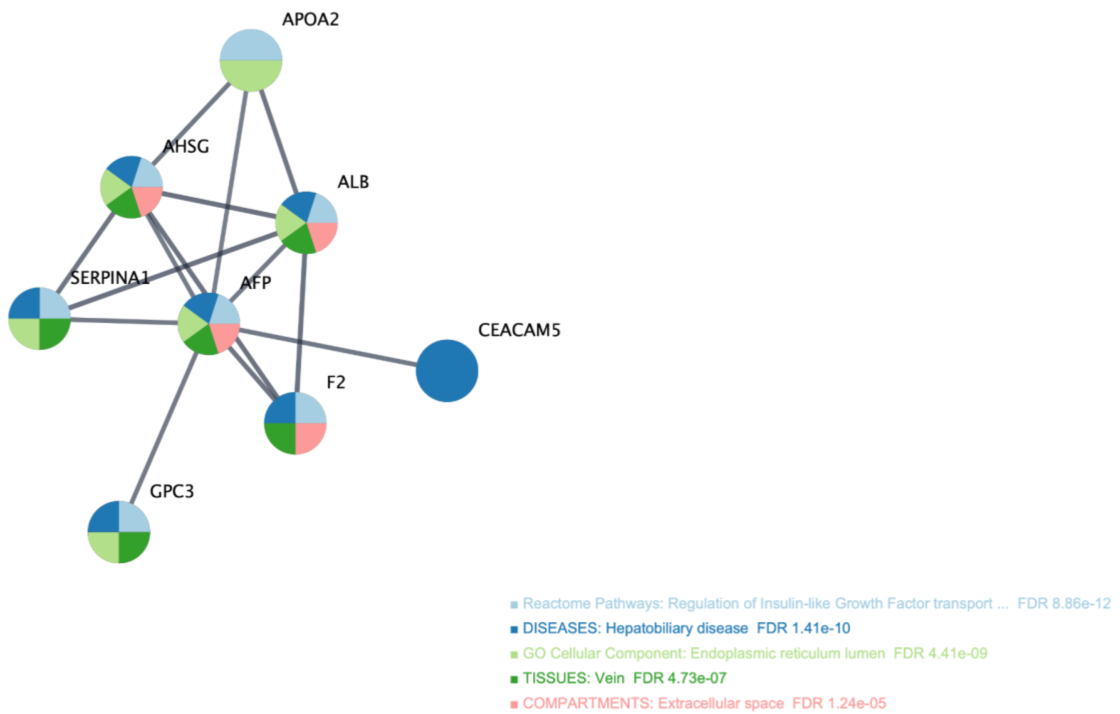


**Supplementary Figure 15. STRING ego network for AFP.**

Nodes are proteins and edges are STRING functional associations in *Homo sapiens* at highest confidence (combined score ≥ 0.90). Coloured pie wedges on each node indicate membership in up to five enrichment terms selected to be non-redundant (Jaccard similarity ≤ 0.9, FDR ≤ 0.05), with each colour representing a distinct enrichment term as indicated in the legend. Link to interactive network in STRING: https://version-12-0.string-db.org/cgi/network?networkId=br27YUaARYHC

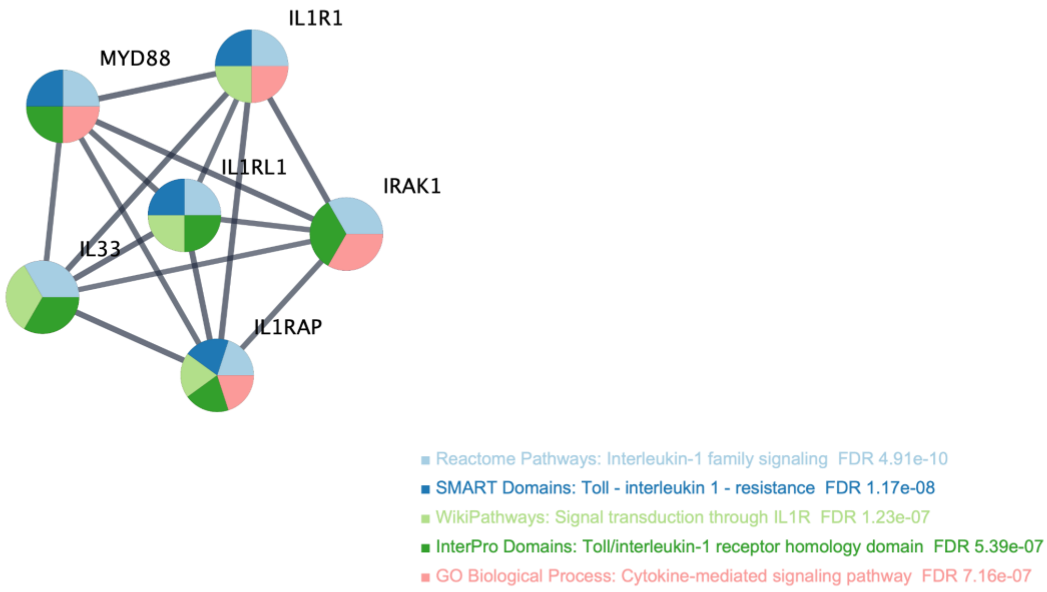


**Supplementary Figure 16. STRING ego network for IL1RL1.**

Nodes are proteins and edges are STRING functional associations in *Homo sapiens* at highest confidence (combined score ≥ 0.90). Coloured pie wedges on each node indicate membership in up to five enrichment terms selected to be non-redundant (Jaccard similarity ≤ 0.9, FDR ≤ 0.05), with each colour representing a distinct enrichment term as indicated in the legend. Link to interactive network in STRING: https://version-12-0.string-db.org/cgi/network?networkId=bDFy3GlAtbew

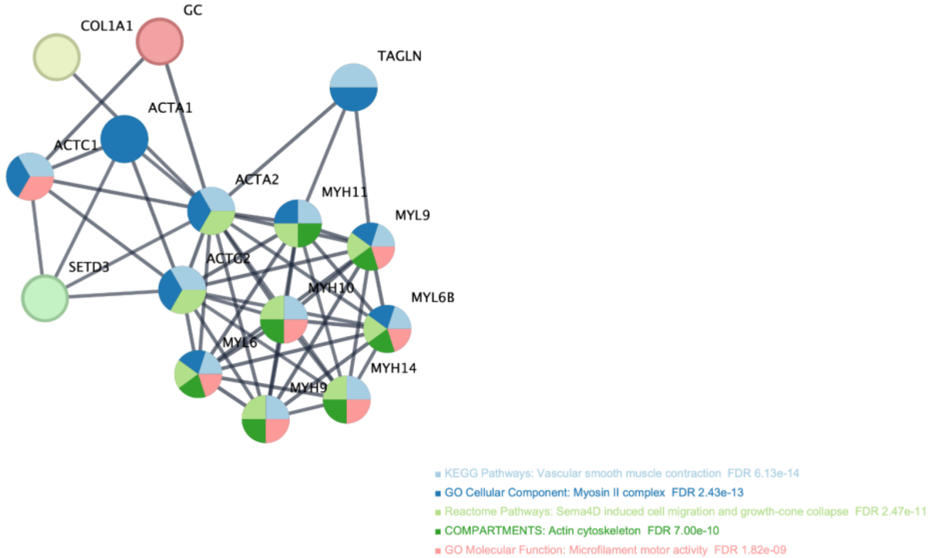


**Supplementary Figure 17. STRING ego network for ACTA2.**

Nodes are proteins and edges are STRING functional associations in *Homo sapiens* at highest confidence (combined score ≥ 0.90). Coloured pie wedges on each node indicate membership in up to five enrichment terms selected to be non-redundant (Jaccard similarity ≤ 0.9, FDR ≤ 0.05), with each colour representing a distinct enrichment term as indicated in the legend. Link to interactive network in STRING: https://version-12-0.string-db.org/cgi/network?networkId=bNML5UX2srDo

The disease association with intestinal pseudo-obstruction (FDR=0.0011) and phenotype association with fetal megacystis (FDR=0.004) enriched in the LMOD1 STRING ego network, provide additional biological context that strengthens our interpretation of LMOD1's role. Both conditions are caused, among others, by loss-of-function mutations in LMOD1 that impair smooth muscle contractility, resulting in peristalsis in the gut and bladder respectively[42,43]. The pathological consequences of LMOD1 absence are the precise inverse of what we propose rising plasma LMOD1 reflects: progressive assembly and activation of the smooth muscle contractile apparatus in the myometrium as delivery approaches. That the same protein whose loss disables smooth muscle contraction should rise in circulation in the weeks before a smooth muscle-dependent physiological event, is internally consistent and supports LMOD1 as a specific marker of smooth muscle activation state rather than a non-specific signal of tissue damage or turnover.

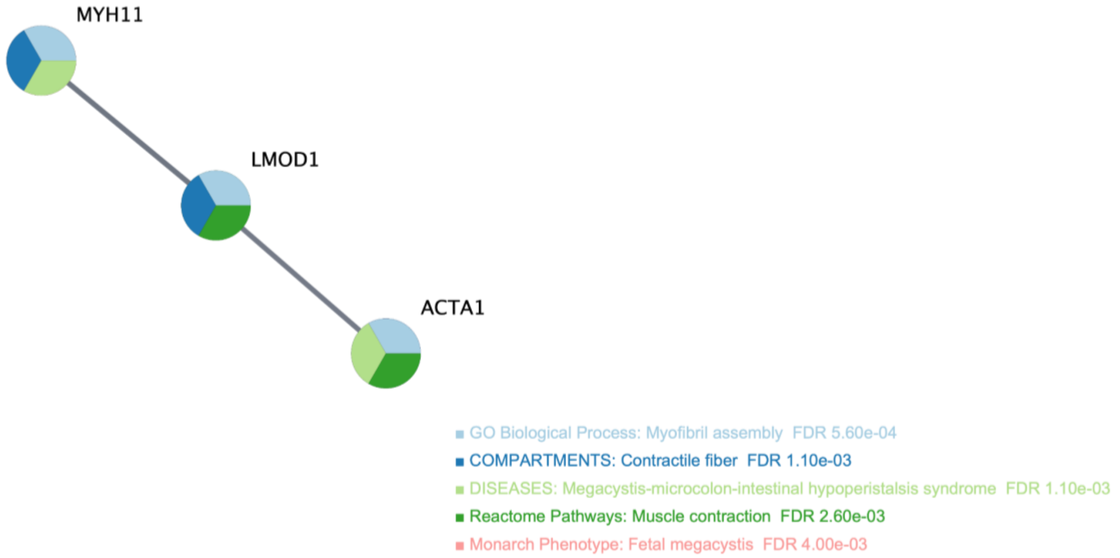


**Supplementary Figure 18. STRING ego network for LMOD1.**

Nodes are proteins and edges are STRING functional associations in *Homo sapiens* at highest confidence (combined score ≥ 0.90). Coloured pie wedges on each node indicate membership in up to five enrichment terms selected to be non-redundant (Jaccard similarity ≤ 0.9, FDR ≤ 0.05), with each colour representing a distinct enrichment term as indicated in the legend. *LMOD1 interactors ACTA1 and MYH11 are also ACTA2 interactors (**Supplementary Figure 17**).* Link to interactive network in STRING: https://version-12-0.string-db.org/cgi/network?networkId=bORDfQFHY0o0

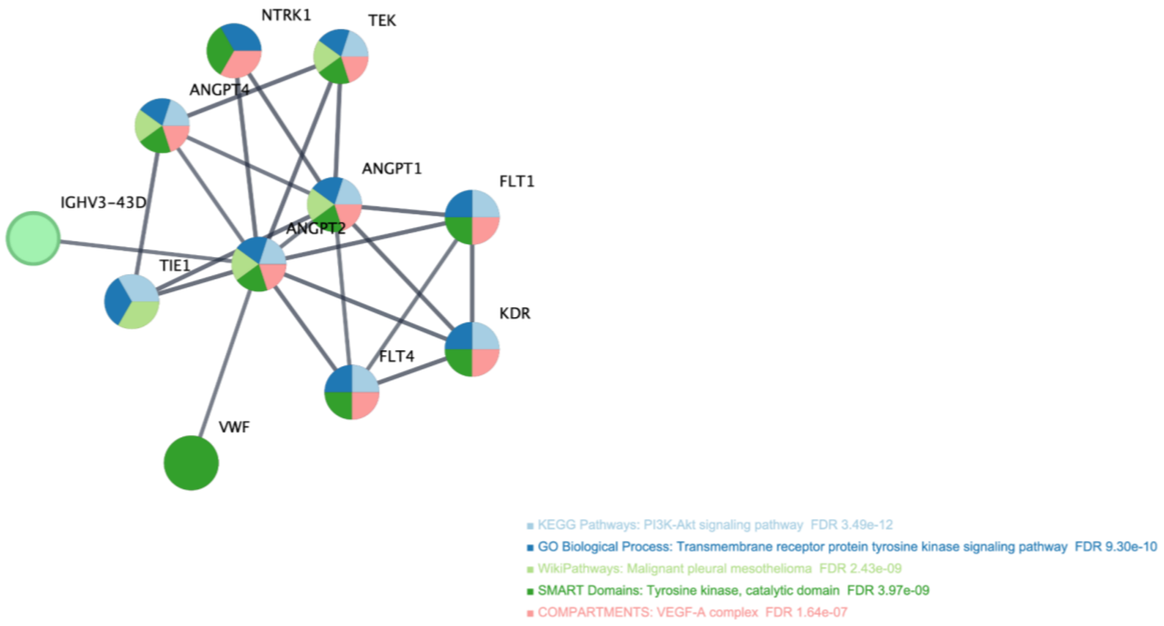


**Supplementary Figure 19. STRING ego network for ANGPT2.**

Nodes are proteins and edges are STRING functional associations in *Homo sapiens* at highest confidence (combined score ≥ 0.90). Coloured pie wedges on each node indicate membership in up to five enrichment terms selected to be non-redundant (Jaccard similarity ≤ 0.9, FDR ≤ 0.05), with each colour representing a distinct enrichment term as indicated in the legend. Link to interactive network in STRING: https://version-12-0.string-db.org/cgi/network?networkId=bfjWZHgYaQiE

### Supplementary Note 8: Comparison with prior work and replication dataset analysis on SomaScan 7k data

Our findings replicate and extend previous longitudinal proteomics studies of term labour. The *Stelzer et al.* study[44], which combined proteomics, metabolomics, and immunomics in a larger population using SomaScan 1317-plex, independently identified IL1RL1 and ANGPT2 among the top proteomic predictors of labour timing – a direct two-protein replication of our cross-method signal using a different platform and population. The five-fold rise of IL1RL1 across pregnancy documented in the *Romero et al.* study[45] provides further independent support for the centrality of the IL-33/ST2 axis in the late-gestational proteomic landscape.

We additionally sought replication in the *Tarca et al.* dataset (GSE206454, SomaScan 7K, n=91 normal pregnancies)[46], which samples from early pregnancy through to approximately two days before delivery. When trajectories were anchored to each woman's earliest available sample and aligned to estimated delivery, IL1RL1 showed a clear rising trajectory with 79/91 women (87%) changing in the expected direction ($p<0.001$), and ANGPT2 showed a clear falling trajectory with 72/91 women (79%) in the expected direction ($p<0.001$), independently replicating our two core signals in a threefold larger population on an independent platform (**Supplementary Figure 20**). The consistent identification of IL1RL1 and ANGPT2 across four independent populations, three platforms, and multiple analytical approaches provides unusually strong cross-study validation for these two proteins as robust markers of labour proximity.

AFP and LMOD1 did not replicate directionally in this dataset (24/91, p=1 and 50/91, p=0.11 respectively), and ACTA2 was not measured on the SomaScan 7K platform. For AFP, the non-replication is biologically informative rather than contradictory: AFP in the GSE206454 dataset spans from the first trimester through to two days before delivery, a window across which AFP first rises and then falls – meaning that the direction of change from earliest to last sample depends entirely on when the first sample was drawn. Women sampled early in pregnancy will show a net rise across the window, while those sampled later will show a net fall, producing the near-random directional result observed (24/91). This is consistent with AFP having a biphasic trajectory across pregnancy, with only the late-gestational decline – captured in our study, which samples exclusively in the third trimester – being relevant to labour prediction. For LMOD1, the trajectory in GSE206454 is essentially flat across the full gestational window examined (**Supplementary Figure 20**), with the expected rise visible only in the immediate pre-delivery period. This dataset does not include samples in the final seven days before delivery for most women, precisely the window in which LMOD1 emerges as the dominant predictor in our analysis, explaining why it does not achieve directional significance here. The replication of the upstream cascade signals (IL1RL1, ANGPT2) and non-replication of the late effector signals (AFP, LMOD1) in GSE206454 is therefore consistent with the proposed temporal structure of the cascade, in which the earliest signals have the broadest detectability window while the latest signals require dense sampling in the final days before delivery.

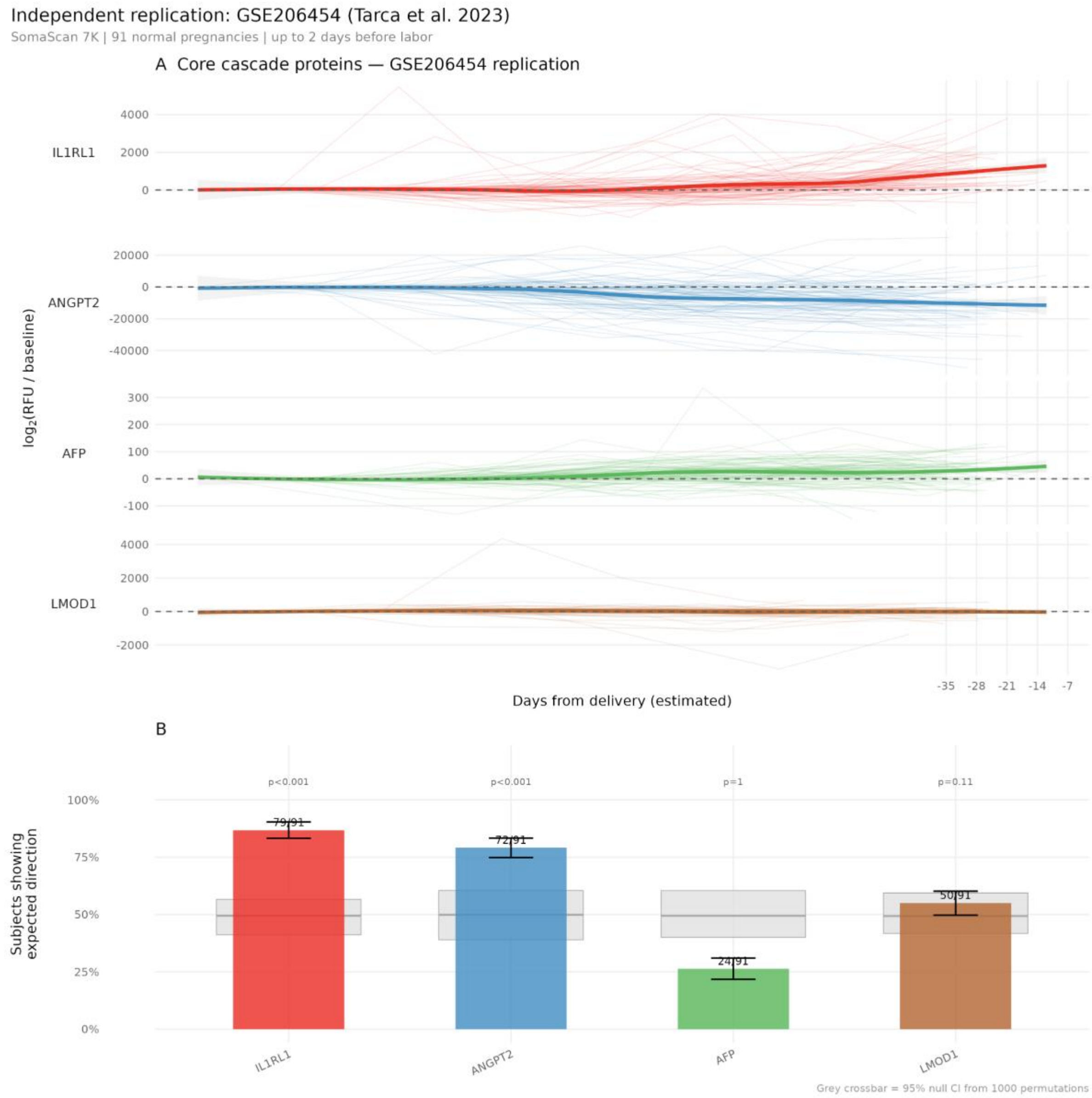


**Supplementary Figure 20. Core cascade proteins in GSE206454 replication dataset.**

(A) Longitudinal trajectories of IL1RL1, ANGPT2, AFP, and LMOD1 in maternal plasma from 91 women in GSE206454, aligned to estimated delivery date. Thin lines represent individual trajectories, and bold lines indicate LOESS-smoothed means with 95% confidence bands. (B) Percentage of subjects showing the expected direction of change (rise for IL1RL1 and LMOD1; fall for ANGPT2 and AFP), with black bars showing observed proportions and grey bars denoting the 95% null CI from 1,000 permutations. P-values assess deviation from the null.

### Supplementary Note 9: Sensitivity analysis of cascade onset ordering

The fixed absolute threshold (|ΔNPX| > 0.8) used to define the day of first detected change (**Figure 3B**, main manuscript) has the theoretical limitation of conflating amplitude and timing: a protein with a large total excursion will cross a fixed threshold earlier than a low-amplitude protein regardless of when it truly begins to change. To assess whether the IL1RL1-first ordering is robust to this methodological choice, we repeated the onset timing analysis using two protein-relative thresholds: 25% and 50% of each woman's own maximum |ΔNPX| for that protein across the antepartum sampling window and compared the resulting cascade ordering to the original fixed-threshold analysis (**Supplementary Figure 21**).

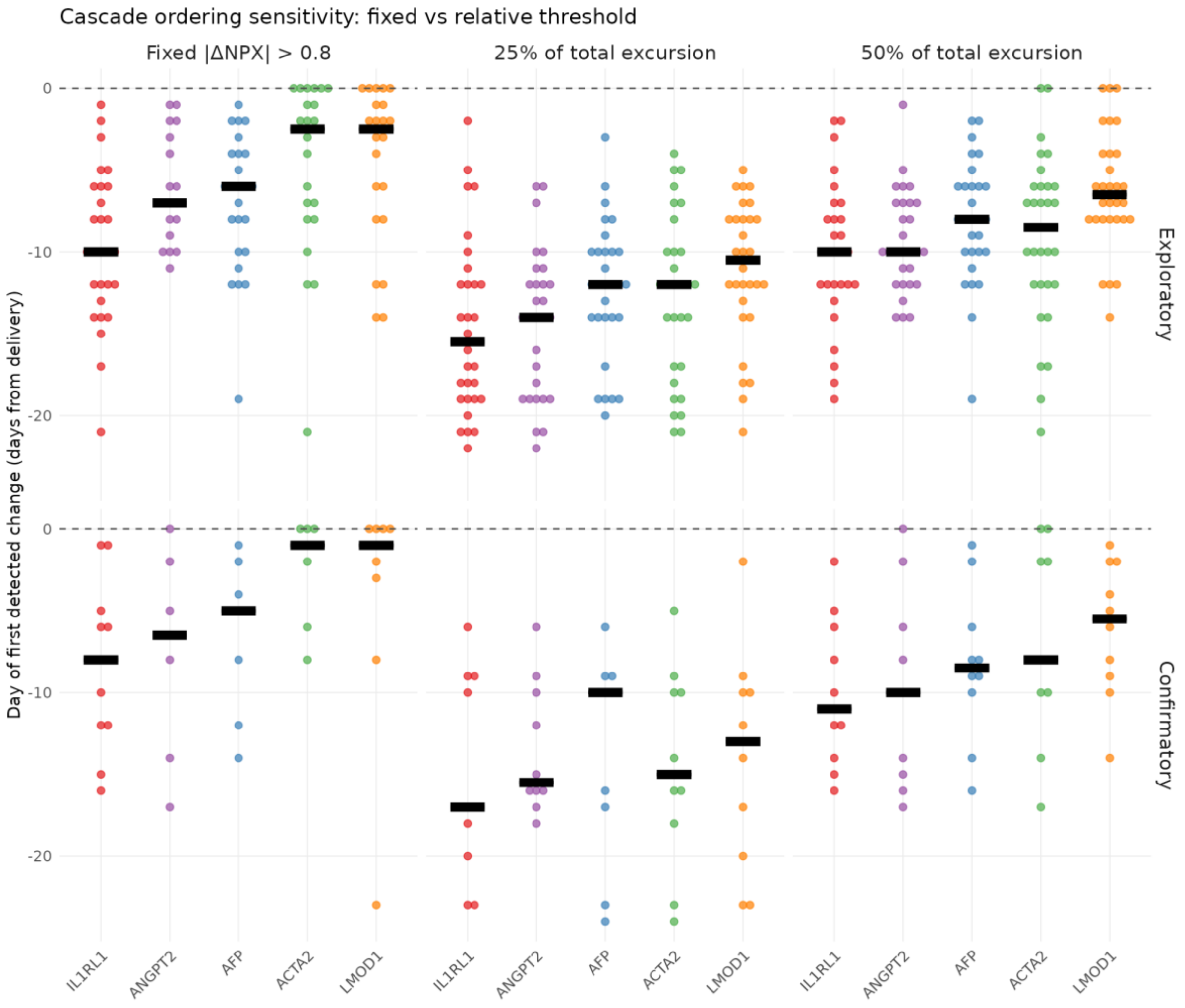


**Supplementary Figure 21. Cascade ordering sensitivity analysis: day of first detected change under three threshold definitions.**

Each column shows onset timing under a different threshold (fixed |ΔNPX| > 0.8; 25% of each woman's total excursion per protein; 50% of each woman's total excursion per protein). Rows show exploratory (n=30) and confirmatory (n=10) groups. Each dot represents one woman; black bars indicate medians. Proteins are ordered along the x-axis according to the hypothesised cascade position. The IL1RL1-first ordering is consistent across all three threshold definitions in both the exploratory and confirmatory groups.

The Kruskal-Wallis test confirmed significant heterogeneity in onset timing across proteins in the exploratory group under all three threshold definitions: fixed threshold (p=0.0005), 25% relative threshold (p=0.017), and 50% relative threshold (p=0.0037). As expected, given the small sample size, no threshold yielded a significant result in the confirmatory group (fixed: p=0.099; 25%: p=0.954; 50%: p=0.354).

### Supplementary Note 10: Mendelian Randomization Analysis

Two-sample Mendelian randomisation was performed for each of the five cascade proteins against three gestational outcomes from the Early Growth Genetics Consortium[47]: gestational duration in days (n=151,987), preterm delivery (n=233,290; 15,419 cases), and post-term delivery (n=131,279; 15,972 cases). Genetic instruments were selected as cis-pQTLs from Loya et al. 2025[48] (UKB Olink Explore 3072, n=47,745), yielding 6 instruments for IL1RL1, 5 for ANGPT2, 2 for AFP, and 1 each for ACTA2 and LMOD1 (**Supplementary Table 6**). No genome-wide significant cis-pQTL was identified for ACTA2 (minimum $p=6.2\times10^{-4}$ in the cis window). Full analytical details are provided in the main Methods section of the manuscript.

Of the five cascade proteins only IL1RL1 yielded a statistically significant MR result (Supplementary Table 7). Genetically higher IL1RL1 levels were associated with shorter gestational duration (β=−0.51 days per SD, 95% CI −0.90 to −0.12, p=0.010, FDR=0.041). There was no evidence of heterogeneity across the six instruments (Cochran's Q=8.18, df=5, p=0.147) and the MR-Egger intercept was near zero (0.016, p=0.903), providing no evidence of directional pleiotropy (**Supplementary Table 8**). The Steiger test confirmed the correct causal direction. Directional effects for preterm (OR=1.038) and post-term (OR=0.945) delivery were consistent with our hypothesis but did not reach statistical significance, likely reflecting lower power for binary outcomes at this effect size.

The direction of the IL1RL1 MR result is consistent with the pathway activation model described in the main manuscript. Genetically higher IL1RL1 levels – reflecting constitutively higher IL-33/ST2 signalling tone – are associated with shorter gestational duration, a direction expected if elevated pathway activity accelerates the parturition programme. This interpretation is supported by the established use of sST2 as a biomarker of IL-33 pathway activity in cardiovascular settings[49], where sST2 rises in proportion to signalling activity rather than as a suppressor of it. We note that the IL1RL1 locus encodes both soluble sST2 and membrane-bound ST2L via alternative splicing, and cis-pQTLs almost certainly perturb expression of both isoforms simultaneously. The MR result therefore reflects the net effect of overall IL1RL1 locus dosage on gestational timing and should be interpreted as evidence for the IL-33/ST2 axis as a whole rather than for the specific contribution of either isoform in isolation.

ANGPT2, and LMOD1 showed null results across all three outcomes, with point estimates close to 0 and no consistent direction. The ANGPT2 null is consistent with a role as downstream hormonal consequence of the progesterone withdrawal cascade rather than a causal driver of delivery timing, as described in the main manuscript. LMOD1 had a single weak instrument and no genome-wide significant cis-pQTL was identified for ACTA2 in our dataset, precluding MR analysis for this protein. ACTA2 is a structural smooth muscle cytoskeletal protein whose circulating levels may be too low or insufficiently heritable for robust pQTL detection in a general population cohort. AFP also showed no significant association with any outcome, and the point estimates did not show a consistent direction across the three outcomes. AFP MR results should be interpreted cautiously as the adult cis-pQTL instrument may not adequately proxy fetal AFP levels in late gestation, since AFP in pregnancy is predominantly of fetal hepatic and yolk sac origin whereas the instrument was derived from non-pregnant adults.

PheWAS of all instrument SNPs against the IEU Open GWAS database ($p<10^{-5}$) revealed distinct patterns by protein (**Supplementary Table 9**). For IL1RL1, non-proteomic associations were confined to eosinophil counts, asthma, atopic dermatitis, and inflammatory bowel disease – all phenotypes within the established IL-33/ST2 biological pathway – consistent with cis-acting effects on receptor expression rather than independent pleiotropy. For AFP, instrument rs3097412 showed strong associations with neutrophil count and CXCL1 levels ($p<10^{-100}$), representing a potential pathway-independent pleiotropic effect that constitutes a second reason, alongside the adult pQTL limitation, to interpret AFP MR results with caution. For ANGPT2, instrument rs2959812 showed associations with heel bone mineral density and diastolic blood pressure. Since the ANGPT2 MR result was null across all outcomes these associations do not affect the primary finding. The single LMOD1 instrument (rs2820323) showed pervasive pleiotropic associations across BMI, body composition, blood pressure, cardiovascular disease, and metabolic traits (>80 associations at $p<10^{-5}$), confirming its exclusion from MR interpretation.

Supplementary Tables 6-9 are provided in a separate excel file (*Supplementary_Tables_6-9.xlsx*) due to their size.

Pregnancy-specific pQTL data at sufficient scale for two-sample MR are not currently available, representing a methodological limitation shared across MR analyses of pregnancy outcomes. For proteins with predominantly adult tissue expression (IL1RL1, ANGPT2, ACTA2, LMOD1), the adult instrument is likely to capture the relevant genetic variation in protein levels, even though pregnancy-specific regulatory effects cannot be excluded. However, as mentioned above, the null result for AFP should be interpreted with caution. Future large-scale proteomics pQTLs studies in pregnant cohorts will be needed to address the limitation and provide more definitive causal evidence.
